# Assembly of the ATP-driven cobalt chelatase

**DOI:** 10.64898/2026.07.21.739949

**Authors:** Yongling Zhou, Hui Yuan, Yongchao Wu, Jian Wang, Huaibo Chen, Liangrui Yao, Mingzhu Wang, Xiao Wang, Jia Wang, Chao He, Xuemin Chen, Lin Liu

## Abstract

Nature has evolved two distinct chelatase families to catalyze the insertion of metal ions into tetrapyrrole macrocycles. Whereas the single-subunit ATP-independent chelatases have been widely investigated, little is known about the three-subunit ATP-driven chelatases. Here we show step-wise assembly of the ATP-driven cobalt chelatase CobSTN that is essential for aerobic vitamin B_12_ biosynthesis. The motor subunit CobS fits into a hexameric or dodecameric spiral, and forms complex with the adaptor subunit CobT. Upon binding to adenine nucleotide, the spiral transforms to an asymmetrical ring and CobT synergistically rotates and inserts a distinctive shaft into the ring hole. The largest subunit CobN interacts with the opposite side of CobT from the CobS ring, and hence the holoenzyme is assembled.

## Introduction

Chelatases catalyze tetrapyrrole metalation with ions, such as iron, magnesium, nickel, and cobalt[1]. The metalated tetrapyrroles either directly produce heme or undergo further modifications to form chlorophyll, coenzyme F_430_, and vitamin B_12_[2]. Based on whether ATP hydrolysis is coupled to the metalation process[3], chelatases are grouped as either the ATP-driven class I chelatases including magnesium chelatase and the cobalt chelatase in aerobic B_12_ biosynthesis pathway, or the ATP-independent class II chelatases including ferrochelatase, nickel chelatase, and the cobalt chelatase in anaerobic B_12_ biosynthesis pathway.

Vitamin B_12_ is synthesized through ∼30 enzymatic steps and only certain microorganisms can accomplish its *de novo* synthesis[4]. Depending on whether oxygen is used and when cobalt is inserted, the steps belong to either the aerobic cobalt-late or anaerobic cobalt-early pathway[5,6]. In the aerobic pathway, class I cobalt chelatase inserts cobalt into hydrogenobyrinic acid *a*,*c*-diamide (HBAD); in the anaerobic pathway, class II cobalt chelatase inserts cobalt into the oxidized precorrin-2. Class I cobalt chelatase is a heterotrimeric enzyme encoded by the *cobS*, *cobT*, and *cobN* genes[7]; class II cobalt chelatase is a single monomeric or homodimeric protein[8]. The latter adopts a bilobal architecture composed of two α/β domains and is possibly evolved from an ancestral chelatase, whose descendants also include ferrochelatase and nickel chelatase[9]. By contrast, the architecture of class I cobalt chelatase remains largely elusive[10,11]. The model of how the CobS, CobT, and CobN subunits are assembled is derived from the magnesium chelatase[12], which is composed of either ChlI, ChlD, and ChlH subunits for chlorophyll biosynthesis or BchI, BchD, and BchH subunits for bacteriochlorophyll biosynthesis[13]. The ATP hydrolytic activity of the motor subunit BchI/ChlI powers magnesium chelation[14]. Crystal structure of *Rhodobacter capsulatus* BchI showed that the AAA+ small subdomain is swapped compared to most AAA+ proteins[15], which serves as a defining structural feature of the clade 7 AAA+ proteins[16–18]. Negative stain electron microscopy (EM) showed that BchI can form a hexameric ring[19], which has also been observed for cyanobacterial ChlI by crystallography and cryo-EM[20,21]. A model of BchI-BchD complex (BchID) has been built from cryo-EM maps[22,23], yet the low resolutions prevented atomic-level insight into the BchID assembly. The adaptor subunit BchD/ChlD links the motor subunit BchI/ChlI with the executor (catalytic) subunit BchH/ChlH[24,25]. The structures of cyanobacterial ChlH were solved by crystallography, which led identification of active site within a cavity[26,27]. However, how the motor, adaptor, and catalytic subunits assemble to accomplish the ATP-driven chelation process is unclear.

Cryo-EM enabled determination of high-resolution structures of ATP-driven machines in the past decade[28]. The ATP-driven chelatases (magnesium chelatase and class I cobalt chelatase) are grouped as members of the clade 7 AAA+ proteins[10,16–18], which are the least characterized ATPases among seven clades of AAA+ superfamily[29,30]. Here, we used single-particle cryo-EM and crystallography to investigate the class I cobalt chelatase from the zoonotic bacterium *Brucella melitensis*[31]. The structures of CobS, CobS-CobT complex (CobST) and the CobS-CobT-CobN holoenzyme (CobSTN) allowed us to capture how the heterotrimeric machine is assembled.

## Results

### CobS homomeric assembly

The 328-residue CobS consists of an N-terminal large AAA+ subdomain (CobS_L_) and a C-terminal small AAA+ subdomain (CobS_S_) (Figs. 1A and 1B). We obtained the recombinant *B. melitensis* CobS protein whose theoretical molecular weight was 37.7 kDa. Size-exclusion chromatography (SEC) showed a broad peak, which suggested that the purified apo-CobS was a mixture of oligomeric states. The purified apo-CobS was subject to cryo-EM analysis, which confirmed that it exists predominantly as hexamer and dodecamer and showed that the dodecamer adopted two major conformations (Fig. S1 and Table S1). However, detailed analysis was limited by the low-resolution maps. In an attempt to optimize the sample for cryo-EM structural analysis, we added ATP to the apo-CobS protein, and obtained the improved density maps for CobS hexamer and dodecamer (Fig. S2 and Table S1). Yet the resolution was still not sufficiently high for model building.

**Fig. 1.**
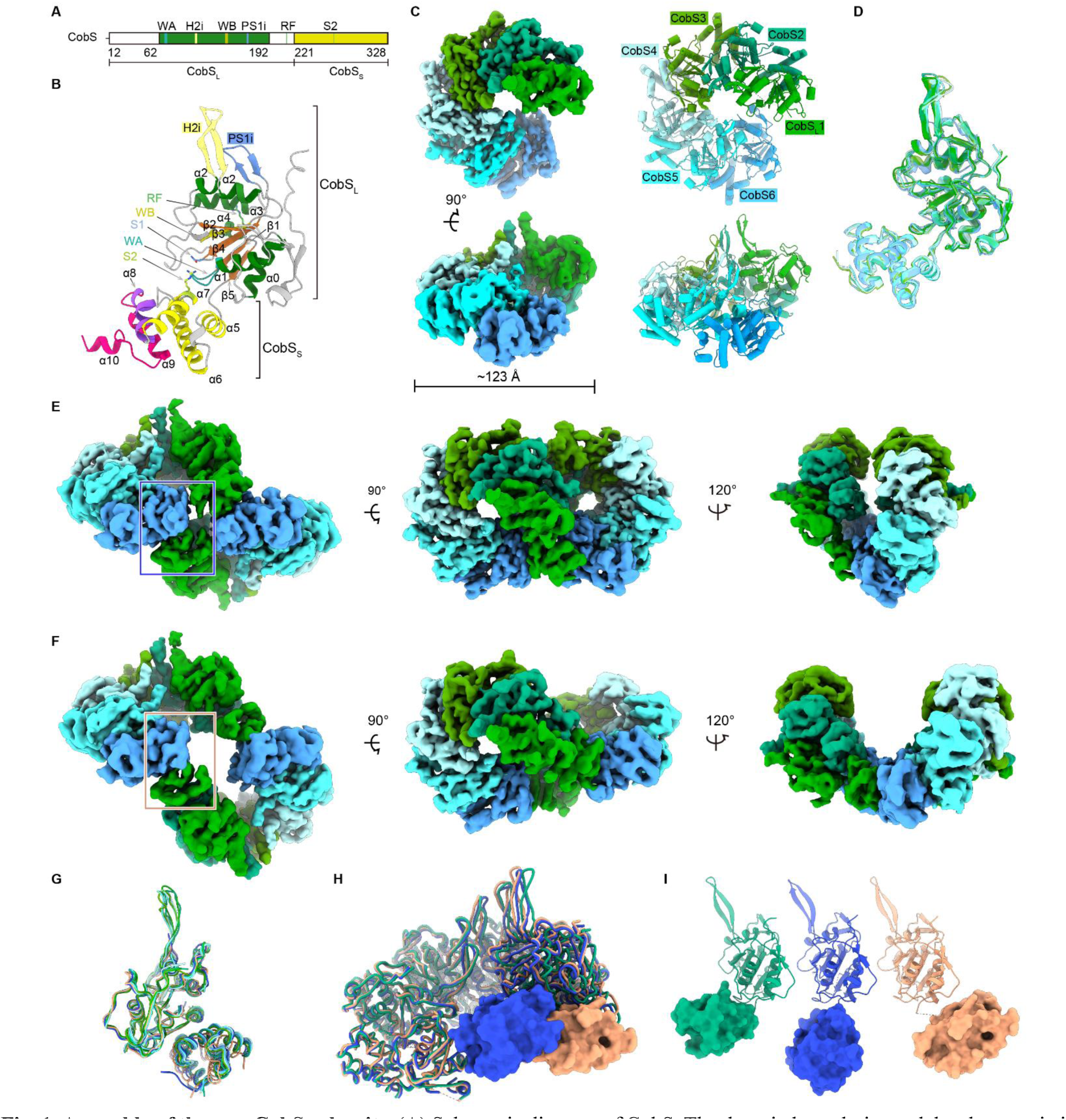
Assembly of the apo-CobS subunits. **(A)** Schematic diagram of CobS. The domain boundaries and the characteristic AAA+ elements are labeled. **(B)** Secondary structures of CobS. α-Helices are labeled and colored by domains as in (A) except the C-terminal helices (α8-α10); β-strands are in brown except those of the AAA+ elements, which are defined and colored as in (A). **(C)** Cryo-EM map and structure of a CobS hexamer. The first CobS_S_ is invisible. **(D)** Superimposition of six CobS subunits. **(E)** Cryo-EM map of the tight conformation of CobS dodecamer. **(F)** Cryo-EM map of the loose conformation of CobS dodecamer. **(G)** *trans*-Interaction between CobS_L_ and its following CobS_S_ (CobS_S_’). The sixth CobS_L_ from one hexamer and the first CobS_S_ from the other hexamer are colored blue and light orange, as indicated by the frames in (E) and (F). **(H)** Superimposition of CobS hexamer. The hexamers from the dodecamers are colored blue and light orange, with the first CobS_S_ in surface representation; the CobS hexamer (C) is colored green. **(I)** Overlay of CobS_L_ to show the swing of CobS_S_. The CobS subunits (*left* to *right*) are the second CobS from (C) and the first CobS from (E) and (F).

The structures of apo-CobS were determined from a heterogeneous sample that contains CobS, the ATP analog adenylyl imidodiphosphate (AMPPNP), and CobT. After 3D reconstruction, the apo-CobS hexamer structure was refined to 3.86 Å (Fig. 1C). Two apo-CobS dodecamer structures and the AMPPNP-bound CobST complex structure were obtained (Fig. S3 and Table S2). Despite being annotated as a clade 7 AAA+ protein[10,17], CobS does not adopt the domain-swapped conformation found in clade 7 proteins. In fact, CobS contains the pre-sensor-1 insert (PS1i) and helix-2 insert (H2i), two specific structural elements for clade 6 and clade 7, but lacks the clade 7-specific pre-sensor-2 insert (PS2i) (Fig. 1B). Both PS1i and H2i are β-hairpins. In the hexamer structure, six CobS subunits arrange into a left-handed spiral and the nucleotide-binding pockets are empty of AMPPNP. Each subunit protrudes a long extension (H2i) surrounding the central channel of the spiral, forming a bulge on the axial end. Therefore, this end is referred to as the convex side; the other axial end, with the channel surrounded by core CobS_L_s and peripheral CobS_S_s, is referred to as the concave side. The six subunits adopt similar structures except that the first CobS_S_ (counted from the convex side) and the sixth H2i extension are missing (Fig. 1D). Along the spiral axis, each CobS subunit slides and rotates about 8.6 Å and 60°, respectively. More CobS subunits could associate with the hexamer from the two axial ends without steric collision, and thus extend the spiral to a tube structure (Fig. S4).

The apo-CobS dodecamer structures were refined to 4.02 Å and 4.21 Å, respectively, for tight and loose states. Both dodecamers can be viewed as a dimer of hexamer exhibiting C2 symmetry (Figs. 1E and 1F). The two bulges point away and the sideview of the dodecamers appears as a V-shape, with the loose state adopting a more relaxed conformation than the tight one. The dimerization interactions occur specifically between the first CobS_S_ and the sixth CobS_L_ domains (Fig. 1G). Interestingly, when compared with interaction between each CobS_L_ and their following CobS_S_, the dimerization interface does not have much difference (Fig. 1G). Despite the significant difference in the overall shape between two dodecamers, the constituting hexamers are nearly the same except the orientation of the first CobS_S_ (Fig. 1H). The difference of dodecamer conformations can be attributed to flexibility of the first CobS_S_, which was unobserved in the apo-CobS hexamer structure (Fig. 1I).

### Apo-CobST assembly

The 637-residue CobT protein has a molecular weight of 70.6 kDa and consists of an N-terminal domain (CobT_N_), a linker of ∼90 residues, and a C-terminal domain (CobT_C_) (Fig. 2A). CobT is predicted to form a two-tiered ring with CobS, similar to a scenario proposed for ChlID[10,15]. We obtained an apo-CobST assembly by co-purification of CobS and CobT, and determined its structure. The 4.24 Å cryo-EM reconstruction showed no such two-tiered ring (Fig. S5 and Table S2). CobS remains as the spiral assembly to which two CobT subunits are attached (Fig. 2B). The first CobS_S_ remained invisible, but an additional CobS_S_ appeared near the sixth CobS_L_, the same position where a CobS_S_ seen in apo-CobS dodecamer structures. One CobT_N_ and two CobT_C_s bind to the concave side, while the linkers are disordered. CobT_N_ peripherally interacts with the fourth CobS_S_, burying 954 Å^2^ surface area (Fig. 2C). Two CobT_C_s are surrounded by the second, third, fifth and sixth CobS subunits, blocking the concave face and totally burying 842 Å^2^ surface area (Fig. 2D). The dimerization interface between two CobT_C_s is of 388 Å^2^, corresponding to only ∼3% of the total surface area of a CobT_C_ (Fig. 2E).

**Fig. 2.**
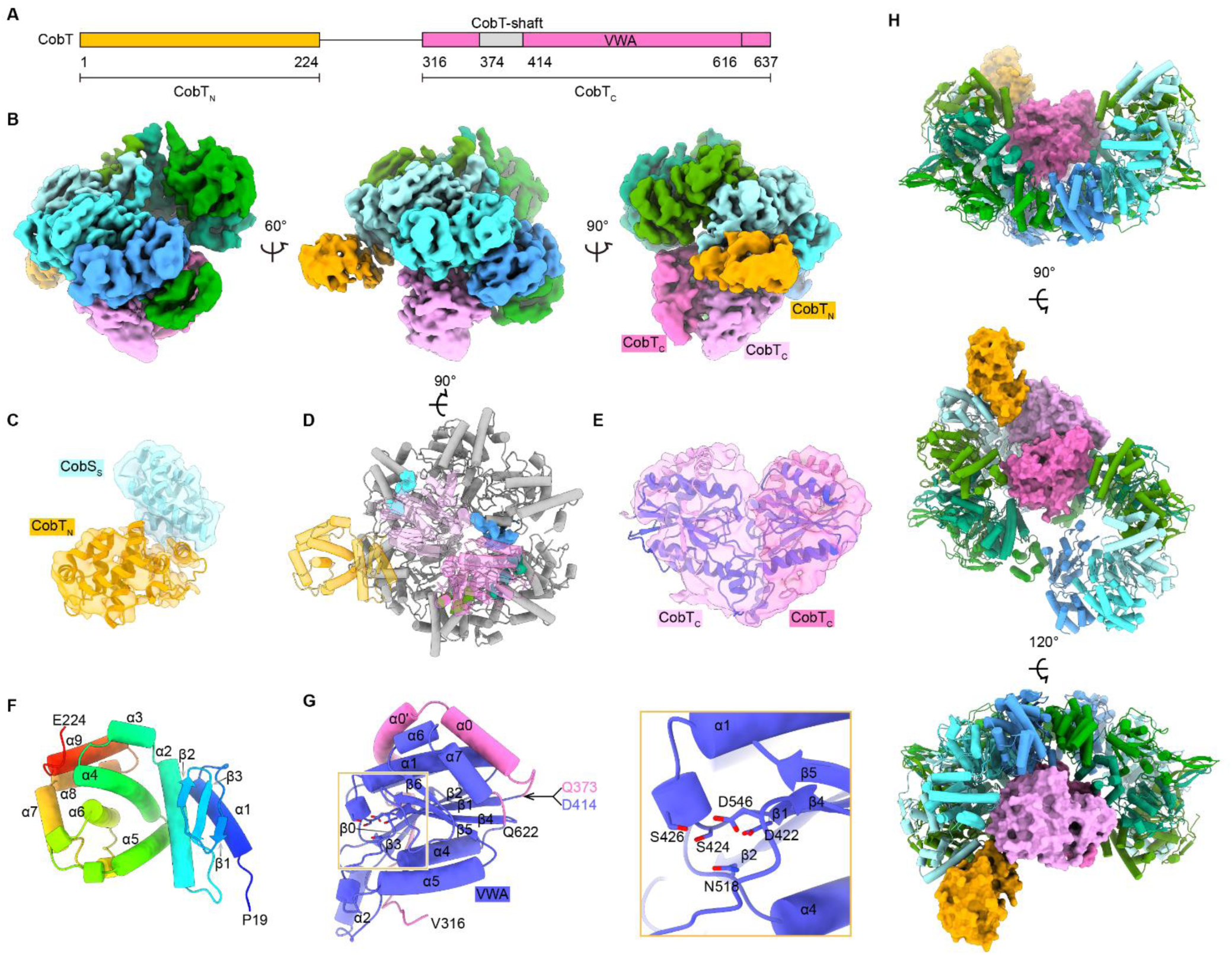
Assembly of the apo-CobST heterocomplex. **(A)** Schematic diagram of CobT. The domain boundaries and shaft fragment are labelled. **(B)** Cryo-EM map of apo-CobST. CobT_N_ is colored orange, and the CobT_C_ dimer is colored pink and light pink. **(C)** Interaction between CobT_N_ and CobS_S_. The density of CobT_N_-CobS_S_ is separated from the overall apo-CobST density. **(D)** Interaction between CobS and the CobT_C_ dimer. CobS (gray) and CobT are in cartoon representation, and the CobT_C_ dimer is in transparency to show the interaction sites. CobS interacting with the CobT_C_ dimer is shown in colored surface. **(E)** Cryo-EM map and structure of CobT_C_ dimer. The density is separated from the overall apo-CobST density. Ribbons from the canonical VWA domain colored are blue, and the other part of the CobT_C_ dimer remain pink and light pink. **(F)** Crystal structure of CobT_N_. The secondary elements are labeled, and the cartoon is colored from blue (N-terminal P19) to red (C-terminal E224). **(G)** Crystal structure of CobT_CΔ_. The secondary elements are labeled, and cartoon is colored as in (E). The linkage corresponding to ΔQ373-D414 is indicated. The inset shows the MIDAS (ball-and-stick) of VWA. **(H)** Fitting apo-CobST into the loose CobS dodecamer.

We then made the CobT_N_ and CobT_C_ truncations and employed crystallography to uncover the structural details of CobT. CobT_N_ crystalized in the presence of the CobS_S_ truncation, and the CobS_S_-CobT_N_ complex crystal was diffracted to 2.2 Å (S3 Table). CobT_N_ contains a three-stranded β-sheet preceded by an α-helix (α1) and followed by eight full α-helices (α2–α9) (Fig. 2F). Structural comparison revealed that CobT_N_ resembled the peptidase M48 subfamily C (M48C) domain. CobT_C_ crystalized when the fragment Q373–R413, which was unobserved in the apo-CobST structure, was deleted. Crystal of this modified CobT_C_ truncation (CobT_CΔ_) was diffracted to 1.48 Å (Fig. 2G and Table S3). Its structure showed a twisted β-sheet sandwiched by α-helices, resembling a von Willebrand factor type A (VWA) domain[32]. CobT shaft sequentially precedes the typical VWA domain which contains a six-stranded β-sheet in the order of 321456, and it is preceded by two tandem helices (α0’-α0). At the very N-terminal end of CobT_C_ lies a short β-strand (β0) parallel to β3 of the VWA β-sheet. Thus, the typical VWA domain in CobT_C_ is half-wrapped by the β0–α0’-α0 segment, and CobT_C_ can be described as a variant of VWA domain with an N-terminal extension. The VWA domain commonly contains a metal ion-dependent adhesion site (MIDAS) for divalent cation. The CobT MIDAS motif is axially distant from the shaft end, situated at the end of β1 and composed of D422, S424, S426, N518, and D546. To visualize how CobT binds to apo-CobS, we compared the CobS spiral of apo-CobST with the apo-CobS dodeacmer assembly (Fig. 2H). A superimposition showed that the spiral overlapped well with the loose dodecamer. No steric collision occurs when the CobT components (one CobT_N_ and two CobT_C_s) are placed into the dodecamer central opening.

### Adenine nucleotide-bound CobS ring

After unsuccessful screening of nucleotide-bound CobS assembly using ATP and AMPPNP for cryo-EM analysis, we generated a defective CobS mutant (CobS^m^), in which the catalytically essential glutamate in Walker B motif (E142) was replaced by a glutamine. The loss of ATP hydrolytic activity of CobS^m^ was experimentally confirmed (Fig. S6). Purified CobS^m^ was mixed with excess ATP and its cryo-EM map was processed to 3.44 Å resolution (Fig. S7 and Table S2). The overall shape of the ATP-bound CobS^m^ structure was a hexamer ring with a diameter of ∼121 Å (Fig. 3A), similar to the shapes of cyanobacterial ChlI[20,21]. A seam appeared on the convex side, making the hexamer an imperfect six-fold symmetry. The subunits were numbered anti-clockwise from the seam, in accordance with the numbering of spiral CobS.

**Fig. 3.**
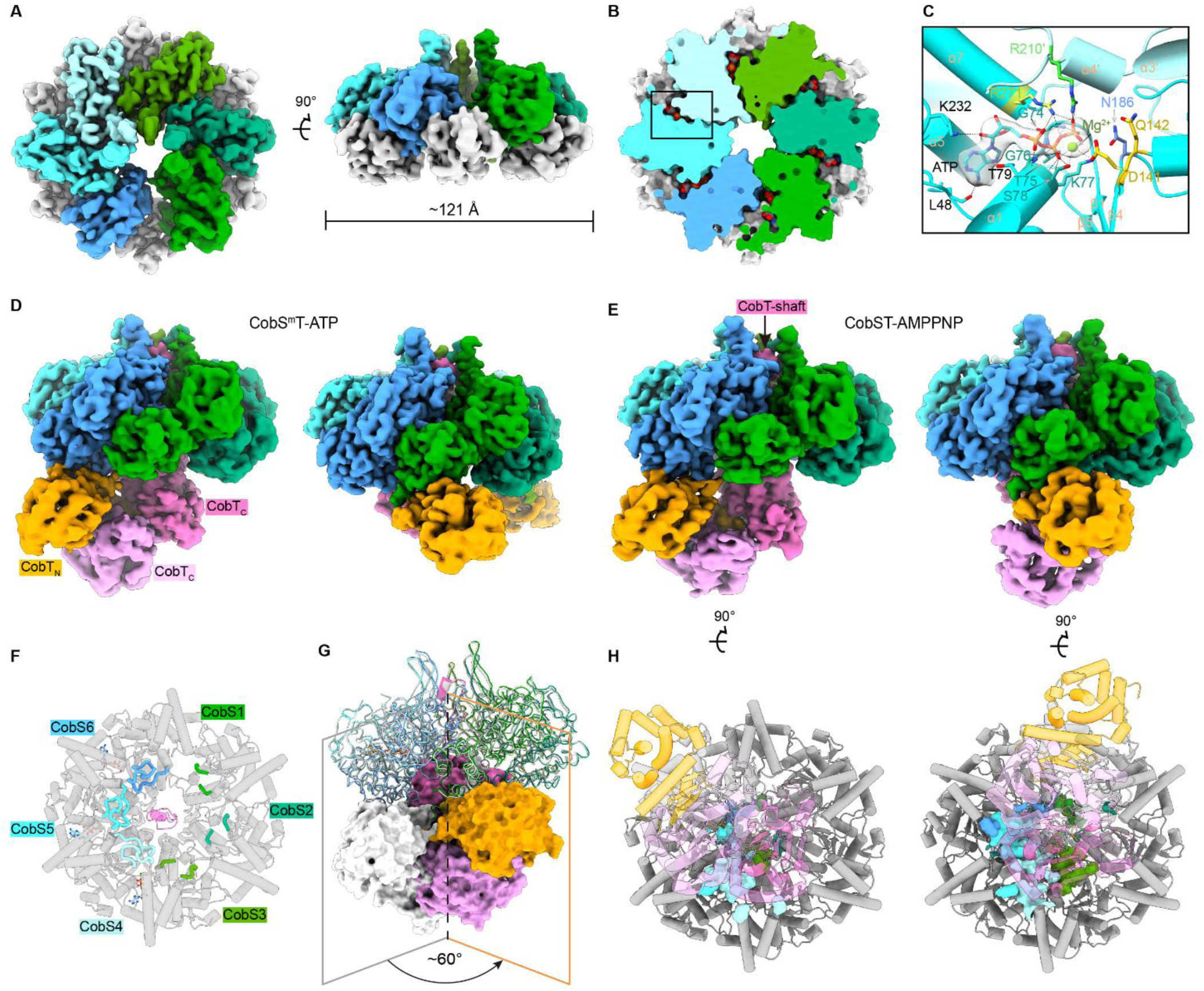
Structure of CobS^m^-ATP and adenine nucleotide-bound CobST. **(A)** Top and side views of cryo-EM map of CobS^m^-ATP. Six CobS_L_ are colored as in Fig. 1C with CobS_S_ in gray. **(B)** Six ATP molecules in the CobS^m^-ATP structure. CobS^m^ is shown as surface, and ATP molecules are in sphere representation. **(C)** Details of ATP binding. The fourth CobS_L_ is colored light blue, and the fifth CobS_S_ is colored cyan. The ATP density is shown in gray. **(D)** Density map of two conformations of ATP-bound CobS^m^T. The color scheme is the same as in (Fig. 2B). **(E)** Cryo-EM maps of two AMPPNP-bound CobST structures. **(F)** Three AMPPNP molecules in the CobST-AMPPNP structure. For clarity, only CobT-shaft is shown while other parts of CobT are omitted. The sensor 1 (N186)-containing loop is shown as thick tube and colored as in (E). **(G)** CobT rotation between the two CobST-AMPPNP structures by superimposing the CobS ring. The CobS hexamers together with CobT-shaft are aligned and shown as ribbons. One conformation is colored gray. **(H)** Interaction between CobS and CobT_C_ dimer in the CobST-AMPPNP structures.

Each CobS^m^ has an ATP molecule bound to its nucleotide-binding pocket (Fig. 3B). The pocket was composed by the Walker A (G71–S78) and Walker B (D141–E142) motifs, sensor-1 (N186), R-finger (R210), and sensor-2 (R271) (Fig. 3C). The *tran*-acting residue R210 contacts the γ-phosphoryl group of the ATP bound to next subunit, except that in the sixth CobS, it is distant from the ATP bound to the first CobS due to the seam.

### Adenine nucleotide-bound CobST assembly

We obtained the ATP-bound CobS^m^T complex and solved its cryo-EM structure, in which all nucleotide-binding pockets in six CobS^m^ were occupied by ATP (Fig. 3D, Fig. S8, and Table S4). While the CobS^m^T-ATP complex appears as an ATP-bound CobS^m^ hexamer ring supplemented with individual CobT components (CobT_N_ and CobT_C_s), substantial conformational changes occur with CobT_C_. The fragment Q373–R413 unobserved in apo-CobST becomes visible and inserts into the central channel of the ring, resembling the central shaft within a motor ring.

We also obtained the AMPPNP-bound CobST complex and solved its structure (Fig. 3E, Fig. S9, and Table S5). One of the two AMPPNP-bound CobST structures is very similar to one conformation of ATP-bound CobS^m^T. However, in AMPPNP-bound CobST structures only the nucleotide-binding pockets of the fourth, fifth, and sixth CobS were occupied by AMPPNP (Fig. 3F). Correspondingly, sensor-1 (N186) loop, part of the nucleotide-binding pocket, is visible only in the fourth to sixth AMPPNP-bound CobS.

When superimposing the CobS hexameric rings from the two conformations, CobT_C_ dimer and CobT_N_ domain rotate together about ∼60° along the axis of motor ring. This means that CobT_N_ can move across the seam, from the sixth to the first CobS_S_, or vice versa (Fig. 3G). Correspondently, the interface between CobT and CobS undergoes a change (Fig. 3H).

### CobS-CobT interactions

ATP-bound CobS^m^T and AMPPNP-bound CobST structures were very similar with respect to CobT_C_ binding. To focus on the CobS ring and CobT-shaft, a local-refined 3.08 Å map was generated for model building. The refined structure provided detailed information on the CobS ring interacting with CobT-shaft (Fig. 4A). Six H2is (long β-hairpins) encircle CobT-shaft, and H2is in turn are surrounded by PS1is (short β-hairpins). The end of CobT-shaft has a short α-helix adjacent to H2is from the third to sixth CobS subunits (Fig. 4B). The polar residues, S102, D110, and R147, that interacts with CobT-shaft, are located in CobS_L_, specifically, on α2, H2i, and α3. Hydrophobic and charged residues are noncontinuously distributed on the shaft, with its end (including a short α-helix) being more hydrophobic. The distal and proximal CobT_C_s forms a similar dimer as in the apo-CobST complex, but in the adenine nucleotide-bound CobST structures, the CobT_C_ dimer rotates so as to insert one CobT shaft into CobS hexamer (Fig. 4C).

**Fig. 4.**
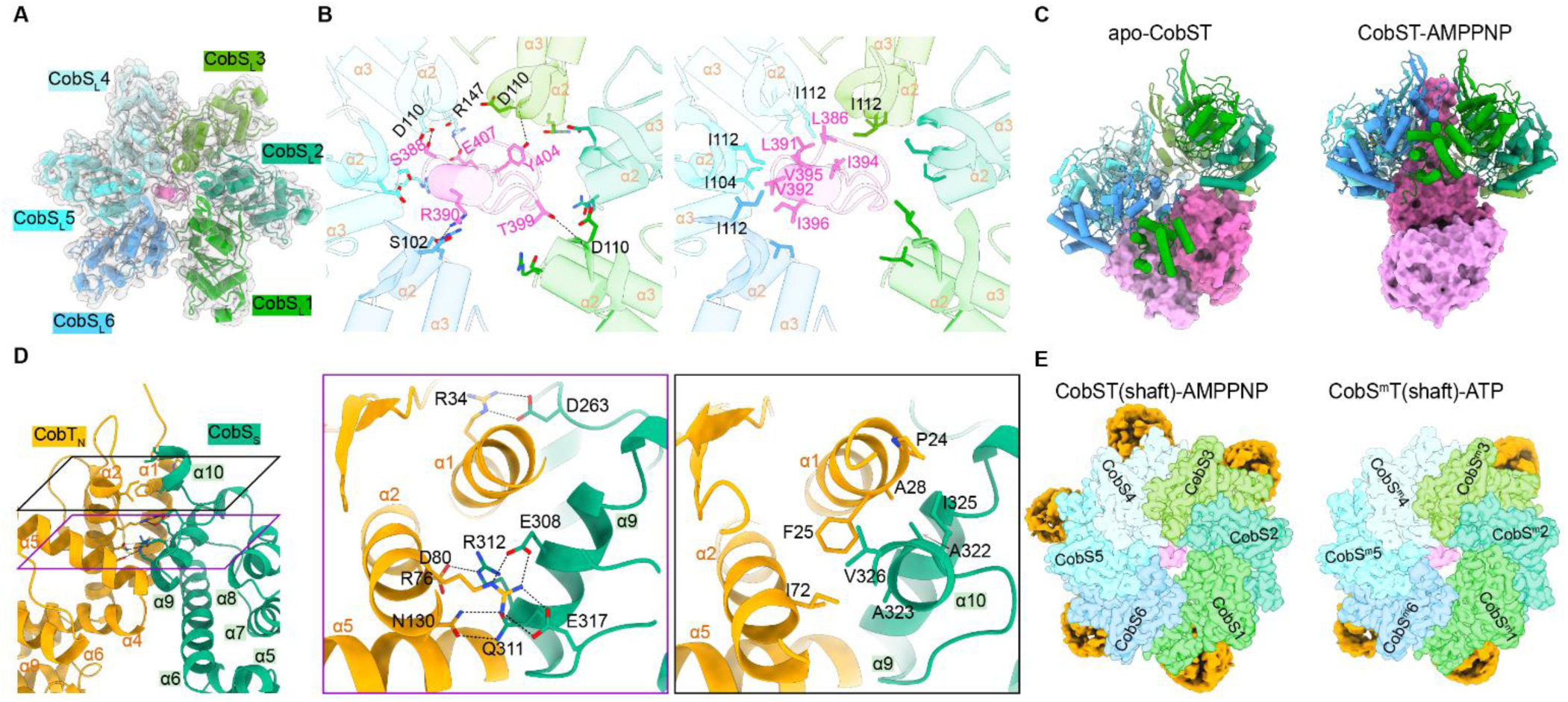
CobS-CobT interactions. **(A)** Interaction between CobS_L_ and CobT-shaft. CobS_S_ is omitted. **(B)** Details of the CobS-CobT-shaft interaction. (*left*) S102, D110, and R147 on each CobS_L_, and the polar residues on CobT-shaft are in ball-and-stick representation. (*right*) I104 and I112 on each CobS_L_, and the hydrophobic residues on CobT-shaft are shown in ball-and-stick representation. **(C)** Rotation difference of CobT_C_ dimer in apo**-**CobST and CobST-AMPPNP. CobT_N_ is omitted for clarity. The two structures are aligned by the first CobS. **(D)** Details of the interaction between CobS_S_ and CobT_N_. Clipped views are zoomed in. **(E)** Possible CobT_N_ binding positions at CobS hexamer. The cryo-EM maps of CobT-shaft, adenine nucleotide-bound CobS (*left*) and CobS^m^ (*right*) are shown, and the other parts of CobT are omitted. The CobT_N_ maps are colored orange. For visualization, the CobT_N_ maps’ level is 1/100 of the CobS map.

In all the available complex structures, CobT_N_ (M48C domain) is seen mainly contacting with CobS_S_ and their interface is conserved in both apo- and nucleotide-bound CobST. We utilized the 2.2 Å CobS_S_-CobT_N_ complex crystal structure for detailed inspection (Fig. 4D). CobS_S_ consists of six α-helices (α5–α10), with α5–α7 belonging to the canonical small helical bundle while α8–α10 corresponding to three extra helices appended to the small bundle (Fig. 1B). Hydrophobic and polar interactions stabilize the complex. Charged residues, including D263, E308, R312, and E317 of CobS_S_, form salt bridges with R34, R76, and D80 of CobT_N_; Q311 of CobS_S_ forms two hydrogen bonds with N130 on α2 of M48C domain. The hydrophobic patch, A322, A323, I325, and V326, on α10 of CobS_S_ form interaction with P24, F25, A28, on α1 and I72 on α2 of M48C domain (Fig. 4D).

We did not observe the linker between CobT_N_ and CobT_C_, consistent with its disordered nature. Lacking of this region in electron density map did hinder the allocation of CobT_C_ to its *cis* CobT_N_. Multiple CobT_N_-binding positions can be seen in previous shown structures, among which the second CobS_S_ offers the least possible position (Fig. 4E). The most often position is next to the seam, i.e., the first or sixth CobS_S_. Simultaneous binding of two CobT_N_s to CobS ring has never been observed in adjacent positions, suggesting that the maximum binding capacity by a CobS ring is three CobTs. An ideal CobST assembly (in 2:1 ratio) could be in three-fold symmetry, but such a structure has not been found and possibly does not exist (Figs. S3A and S10A).

### CobSTN holoenzyme

To explore the assembly of CoSTN holoenzyme, we expressed and purified the 137-kDa CobN. The holoenzyme was obtained by mixing CobN with the AMPPNP-bound CobST in a 5:1 ratio in the presence of Co^2+^ ion. We were able to resolve the AMPPNP-bound holoenzyme structures at ∼4.2 Å resolution and the CobN structure at ∼3.5 Å resolution (Fig. S10 and Table S6). The CobSTN holoenzyme can be described as a three-tier architecture (Fig. 5A). Six CobS ATPase subunits constitute the bottom tier, a CobT subunit serves as the middle tier, and a CobN subunit sits on top.

**Fig. 5.**
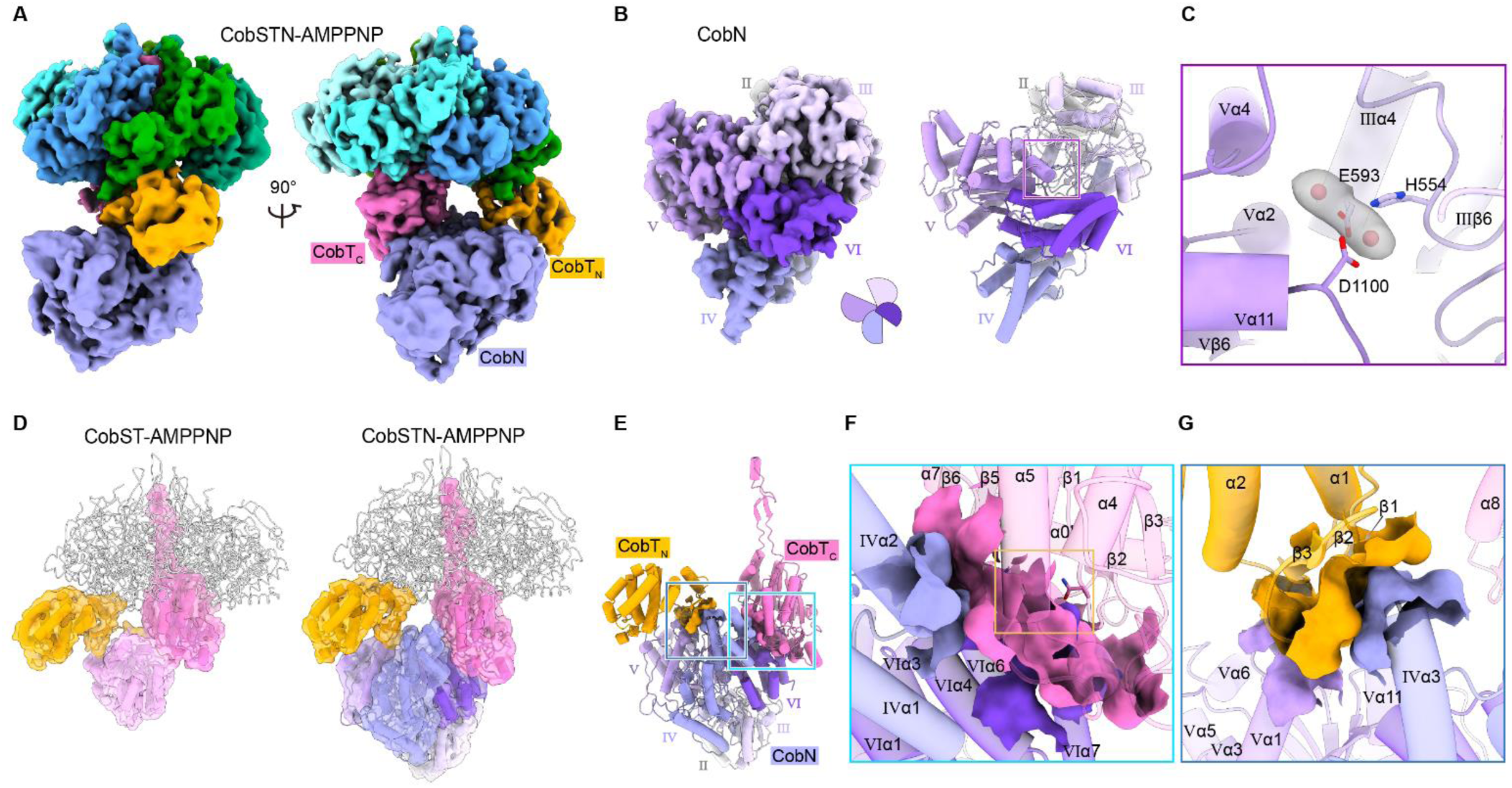
Structure of the CobSTN holoenzyme. **(A)** Cryo-EM map of CobSTN holoenzyme with AMPPNP molecules. CobN is colored purple. **(B)** Map and structure of CobN. The pliers sketch mimics the CobN domains. **(C)** Two Co^2+^ ions in the proposed active site between domains Ⅲ and Ⅴ. Co^2+^ ion is shown as red sphere, with its maps colored gray. **(D)** Comparison of the CobST-AMPPNP and CobSTN-AMPPNP structures. The aligned CobS hexamers are shown as gray ribbons. **(E)** Interaction between CobN and CobT. **(F)** Zoom-in view of CobT_C_-CobN interaction. The secondary structures are labelled, and the position of VWA MIDAS is indicated by an orange box. **(G)** Zoom-in view of CobT_N_-CobN interaction.

Previous crystal structure of *Mycobacterium tuberculosis* CobN has revealed that this large subunit can be divided into six domains (I–VI)[11]. Four domains (I–III and V) are αβα-sandwiches, and the other two domains (IV and VI) are mainly helical bundles. Domains I and II appear as the head and neck to the body assembled by domains III–VI. The cryo-EM structure of CobN reported here contains domains II–VI, with the head domain lacking electron density (Fig. 5B and Fig. S11A). Two Co^2+^ ions are found within the putative active site composed of putatively critical residues including H554, E593, and D1100 (Fig. 5C). To form a holoenzyme, CobN replaces the distal CobT_C_ (Fig. 5D). Resembling the armadillo-repeat α-solenoid that often serves a platform for protein-protein interaction[33], domain VI provides the largest CobT-interacting surface (Fig. 5E). The last two α-helices of domain VI and their connecting loop locate very close to CobT MIDAS, implying a possible role of MIDAS on metal-related function. The lobe-shaped domain IV provides a small CobT_C_-interacting surface by interacting with the VWA, far from the long bent α0’-α0 (Fig. 5F). Furthermore, domain IV interacts with CobT_N_, specifically the three-stranded β-sheet of the M48C domain; domain V interacts with the β3-α2 loop of CobT_N_ via its α1-β1 loop (Fig. 5G). CobN sits between the N- and C-terminal domains of CobT, without touching any CobS subunits.

The holoenzyme structure allows us to assess the assignment of CobN domains. Its six domains were defined according to sequence similarity to ChlH[11,26] (Figs. S11B-S11G), when no further structural information was available then and we were not able to assign a function to any domains. The head and neck domains (I and II) are rightfully named with respect to the three-tier CobSTN assembly. The two large αβα-sandwich domains (III and V) encompass the putative active site, with each having an appendage: the lobe domain IV (to III), and the α-solenoid domain VI (to V). To simplify, the four body domains (III–VI) can be analogized as a pair of pliers (Fig. 5B). The lobe and α-solenoid domains act as two handles, and the two large αβα-sandwich domains are the jaw responsible for pressuring the substrates, Co^2+^ and HBAD.

## Discussion

The structures suggest an assembly model starting from individual CobS, CobT, and CobN subunits (Fig. 6). CobS lacks PS2i and does not adopt a domain-swapped structure seen in BchI/ChlI[15,20,21]. This clearly shows that class I cobalt chelatase belongs to clade 6 instead of clade 7 AAA+ proteins to which magnesium chelatase belongs, and is in agreement with that clade 7 could have diverged from clade 6[16]. However, the evolutionary history of chelatases indicates that magnesium chelatase has appeared earlier than class I cobalt chelatase[2,34]. This discrepancy reflects the complexity of class I chelatases, which is also manifested by the trimer-of-dimer arrangement for BchI assembly[23].

**Fig. 6.**
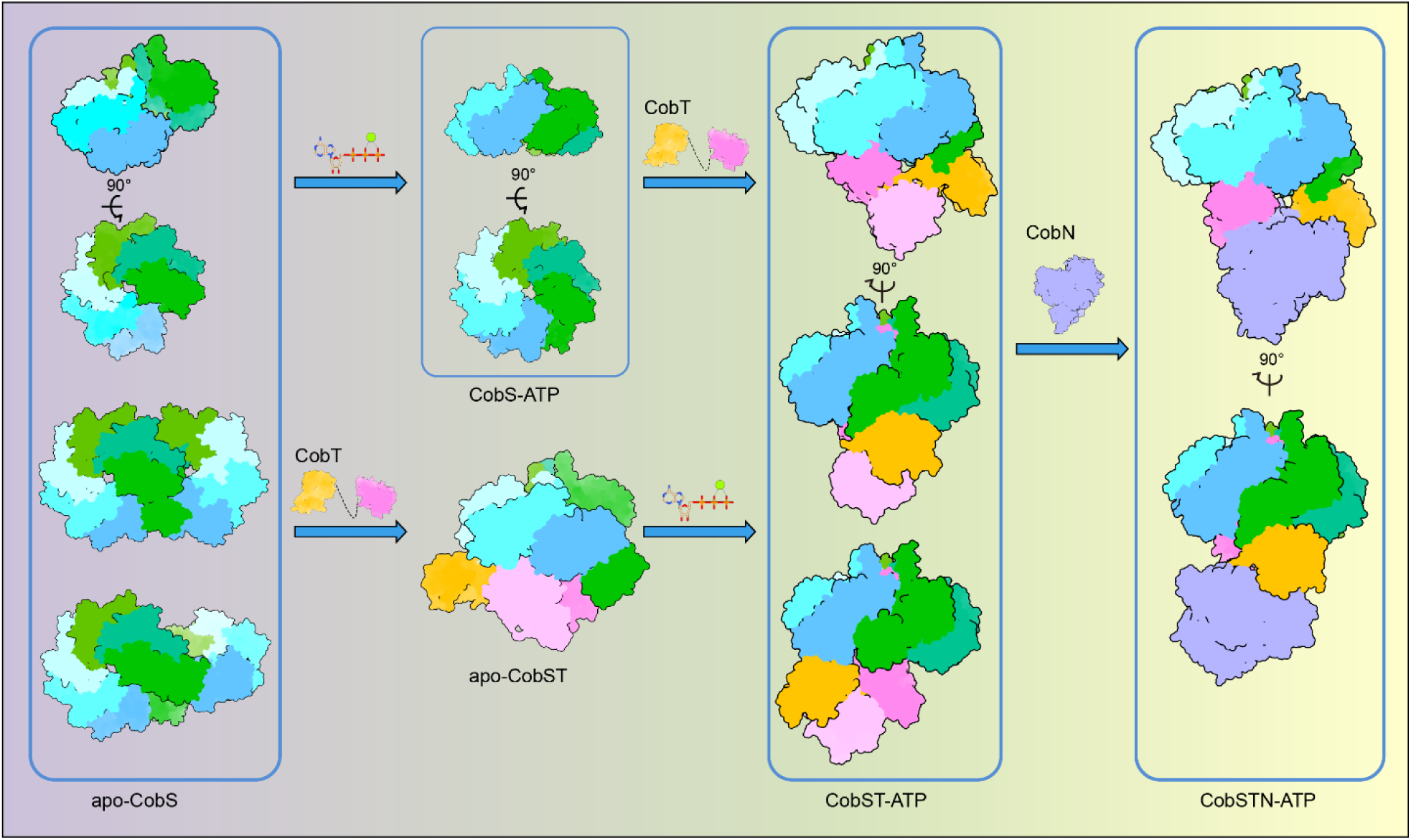
Model of the assembly of CobSTN holoenzyme.

The VWA domain-containing protein associated with AAA+ proteins are annotated as the MoxR family of the AAA+ superfamily[35]. A member of the MoxR family can serve as an adaptor between the effector and motor[36]. Interestingly, the MoxR proteins were initially proposed as molecular chaperones for metal insertion processes[37]. A similar motor-adaptor-effector model has been proposed for iron insertion into cytochrome *c*-dependent nitric oxide reductase (*c*NOR)[38]. The membrane-spanning enzyme *c*NOR is a heterodimer composed of NorB and NorC, with NorB containing two heme irons, a non-heme iron (Fe_B_), and a calcium[39]. Insertion of Fe_B_ into NorB is catalyzed by the ATPase NorQ and the VWA domain-containing NorD[40]. The proposed NorQ-NorD-NorB model, in a molar ratio of 6:1:1, resembles the CobSTN holoenzyme reported here, and thus implicates functional conservation of such an assembly. Carbon monoxide dehydrogenase, coupled to bacterial respiratory chain, utilizes the bimetallic [CuSMoO_2_] cluster for electron transfer[41]. Biogenesis of this metal cluster requires the BchI homolog CoxD and the VWA domain-containing CoxE, yet the metalation process is still unclear[42]. Carbon dioxide fixation depends on the inefficient enzyme Rubisco, whose maintenance entails sets of chaperone-like proteins[43]. CbbQO, a bipartite Rubisco activase composed of the motor CbbQ and adaptor CbbO, catalyzes Rubisco remodeling in chemoautotrophic bacteria. ATP hydrolysis by the CbbQ hexamer provides energy to remodel Rubisco through one CbbO adaptor[44,45]. Regulatory ATPase variant A (RavA) and VWA interacting with AAA+ ATPase (ViaA) constitute a motor-adaptor assembly, while its biological function remains controversial with respect to the effector candidates[46–49].

CobN contains four αβα-sandwich domains that can be regarded as two heterodimers (I-II and III-VI). The directions of β-sheet cores converge to the centers of heterodimers, similar to that of class II chelatases. HBAD could be able to bind to both the I-II interface and III-VI interface. Given the similarity of CobN to ChlH and the active site information of the latter[27], we propose that the III-VI interface encompasses the site for chelating Co^2+^ into HBAD. The I-II interface might be involved in delivery of corrin substrate and product, or other tetrapyrroles. Indeed, in the cryo-EM structure of *Bacteroides thetaiotaomicron* CobN homolog, a heme binds to this interface[50]. This CobN homolog has been suggested to extract iron from heme, and thus is referred to as a heme dechelatase.

## Materials and Methods

### Construction and expression

The *cobS*, *cobT*, and *CobN* genes from *B. melitensis* were codon optimized for expression in *Escherichia coli* (Sango Biotech, Shanghai, China). The synthesized genes were separately cloned into expression plasmids. We also constructed the plasmids encoding CobS^m^, CobS_S_ (P221–A328), CobT_N_ (M1–S226), and CobT_CΔ_ (V316–R637, ΔN374–R413) using the corresponding wild-type templates. The pET constructs were separately transformed into *E. coli* Rosetta2 (DE3) competent cells (Novagen, Beijing, China) for expression. For co-expression of CobS_S_ and CobT_N_, pET-28a(+)-His_6_-cobT_N_ and pET-22b(+)-cobS_S_ were co-transformed into the BL21(DE3) cells. The cells were grown at 37°C in Luria-Bertani medium, and when the optical density at 600 nm reached 0.6, induction was achieved by addition of isopropyl β-D-thiogalactoside to a final concentration of 0.4 mM. The cells were grown at 16°C for additional 16 hours before harvesting by centrifugation (3,000 g). For expression of CobN, the pETMALc-H-MBP-His_6_-TEV-CobN construct was transformed into the Rosetta2 (DE3) cells, and the cells were grown at 37°C in the ZYM-0502 self-induction medium[51]. After the optical density at 600 nm reached 0.6, the cells were grown at 20°C for additional 48 hours before harvesting.

### Purification and complex preparation

The His-tagged proteins (untruncated CobS and CobT) were purified by a two-step process using nickel affinity column and size-exclusion chromatography (SEC). The harvested cells were resuspended in buffer A (150 mM NaCl and 20 mM Tris-HCl, pH 7.5), and sonicated in an ice-water bath. The lysate was centrifuged (13,000 g) and the supernatant was loaded onto an Ni-NTA column (QIAGEN, Shanghai, China), which was equilibrated with buffer A. The column was washed with 50 column volumes of 20 mM imidazole in buffer A, and then the protein of interest was eluted with 20 column volumes of 200 mM imidazole in buffer A. The eluate was concentrated by ultrafiltration (3,000 g) to ca. 2 mL using Amicon Ultra-15 mL Centrifugal Filter Unit, and then loaded onto a HiLoad 16/60 Superose 6 column (Cytiva, Shanghai, China), which was equilibrated and eluted with buffer A. Peak fractions were collected and analyzed by SDS-PAGE. The purification of CobT_CΔ_ was the same as the two-step process except two differences: the buffer was 150 mM NaCl, 10% glycerol, and 20 mM Tris-HCl, pH 7.5. The tag-free CobS^m^ expressed by the pETMALc-H construct was purified by a three-step process, with the first and third steps being same to the two-step process for His-tagged proteins. The second step was the removal of the MBP-His tag by TEV protease. After the first step (nickel affinity column), MBP-His_6_-CobS^m^ was mixed with the His-tagged TEV protease at a mass ratio of 1:20, and the mixture was kept with gentle shaking at 4°C for 10 hours. The mixture was loaded onto a Ni-NTA column and the flow through was collected and concentrated before the third step (SEC). This three-step process was also applicable for CobN purification.

The CobST complex was obtained by co-purification of the tag-free CobS and His-tagged CobT. One liter of CobS-expressing cells were mixed with three liters of CobT-expressing cells before harvesting. The purification process was the same as for the His-tagged proteins. The preparation of ATP or AMPPNP-bound complex was obtained by mixing the purified protein, MgCl_2_, and ATP (or AMPPNP) at a molar ratio of 1:100:100 at 37°C for 5 minutes, and then subjecting to SEC. Buffer A supplemented with 5 mM MgCl_2_ was used for equilibration and elution. The CobS_S_-CobT_N_ complex was obtained by co-expression. Its purification was the same as the two-step process except two differences: the buffer (200 mM NaCl and 20 mM Tris-HCl, pH 7.5) and SEC column.

### Cryo-EM sample preparation and data collection

The protein concentration was adjusted to 1 mg/mL, and all samples were centrifuged at 13,000 g at 4°C for 5 minutes before cryo-EM sample preparation. A 3-µL sample was spotted on a freshly glow-discharged QUANTIFOIL R 1.2/1.3 (Cu, 300 mesh) grids at 22°C, blotted for 3 s, and was stored in liquid nitrogen before data collection. To prepare the CobS^m^-ATP or CobS^m^T-ATP complex, a 20-fold molar excess of ATP was added to the sample and incubated at 37°C for 5 minutes before concentration adjustment. To prepare the AMPPNP-bound CobSTN holocomplex, CobST-AMPPNP, CobN, and CoCl_2_ were mixed at a molar ratio of 1:5:5 and incubated at 37°C for 5 minutes before concentration adjustment. The frozen grids were loaded onto a Glacios electron microscope operated at a 200 kV at the cryo-EM Facility of Anhui University. Automated image acquisition was carried out with EPU software. Movies were recorded in EER format on a Falcon 4 direct electron detector paired with Selectris energy filter using the counted mode at 165K nominal magnification (calibrated pixel size of 0.698 Å) at -0.6 µm to -2.2 µm defocus. During data collection, the total dose was 50 e^-^/A^2^.

### Cryo-EM structure reconstruction and model refinement

Cryo-EM data was analyzed using CryoSPARC[52]. All frames in each collected movie were aligned and summed using Patch Motion Correction, and contrast transfer function (CTF) estimation was performed using Patch CTF Estimation. Blob picker and template picker were used for particle picking. The extracted particles were subjected to 2D classifications and 3D classifications to remove junk particles and to screen for the most homogeneous particles for in-depth 3D structural analyses. The final 3D reconstruction for each class was performed using cryoSPARC non-uniform refinement. The resulting map was post-processed with DeepEMhancer[53] before model building. The resolution was based on the Gold standard refinement procedure and the 0.143 Fourier Shell Correlation (FSC) threshold. Local resolution was calculated using Local Resolution Estimation. Model building was performed with AlphaFold2 model[54] as an initial structure to fit into the maps using UCSF Chimera[55]. The model was then manually adjusted and rebuilt according to the cryo-EM density in Coot[56]. Real-space refinement was done with Phenix[57].

### Crystallization and structure determination

The purified CobS_S_-CobT_N_ complex was concentrated to 16 mg/mL. Crystal screen was performed using sitting-drop vapor diffusion method at 18°C. Crystals appeared within 4 days under the condition of 3.2 M sodium formate and 16% glycerol. The CobT_CΔ_ protein was concentrated to 8 mg/mL and then screened. Crystals appeared within 8 days under the condition of 0.1 M sodium acetate pH 4.5, 25% PEG 3,350, or 0.1 M sodium citrate pH 5.6, 20% PEG 4,000. Before diffraction, crystals were transferred to an antifreeze solution (crystallization condition supplemented with 25% glycerol) and were flashly cooled in liquid nitrogen. X-ray diffraction data were collected at beamlines BL18U1 and BL19U1 of the National Facility for Protein Science in Shanghai. The collected data (Table S2) were processed with the HKL-3000 program package[58]. The structure was solved by molecular replacement with AlphaFold2 model[54] as search template using the CCP4 suite[59]. The structure was iteratively refined with Coot[56] and Phenix[60]. The model quality was accessed by MolProbity[61].

### Assay of ATPase activity by HPLC

AMP (Sigma-Aldrich, Shanghai, Sigma) was dissolved in water with a concentration of 0.25 mM. The samples were centrifuged (13,000 g) for 10 minutes at 4℃ before being subjected to HPLC detection. ADP was dissolved in buffer A, with the concentration and sample preparation conditions being the same as those of AMP. The ATP standard sample was prepared in the same way as ADP. The ADP-1 standard sample was prepared by incubating the sample at 37℃ for 5 minutes, then at 100℃ for 5 minutes, and centrifuged (13,000 g) for 10 minutes at 4℃ before HPLC detection. The preparation conditions for ATP-1 standard sample were the same as those for ADP-1 standard.

Apo-CobS, MgCl_2_, and ATP were mixed in a molar ratio of 1:100:100. The mixture was prepared under the same conditions as the ADP-1 sample, maintaining the final concentration of ATP at 0.25 mM. HPLC detection was performed using Thermo Scientific Hypersil GOLD C18 (250×4.0 mm, 5 μm). The equilibration of the chromatographic column and the elution of the samples were carried out using a 10% A/90% B mobile phase (A: acetonitrile, B: 50 mM NaH_2_PO_4_, 50 mM Na_2_HPO_4_, 10 mM TBAB). 60 μL of standard sample and the samples from the apo-CobS+ATP experimental group were injected into the HPLC instrument (Anhui Wanyi, China). The detection wavelength was 254 nm, the column temperature was 25°C, the sample elution flow rate was 1 mL/min, and the sample elution time was 25 min. The ATP hydrolase activity of the apo-CobS protein was studied by analyzing the HPLC detection results of the apo-CobS+ATP group and the standard sample group.

Apo-CobST, MgCl_2_ and ATP were mixed in a molar ratio of 1:10:10. The ATP hydrolase activity of the apo-CobST complex was studied using the HPLC method for the ATP hydrolase activity of apo-CobS protein.

CobS^m^ was mixed with ATP at a molar ratio of 1:20 (with MgCl_2_ in the buffer of CobS^m^ protein), and then the ATP hydrolase activity of CobS^m^ protein was studied according to the HPLC detection method for CobS protein ATP hydrolase activity. The method for the ATP hydrolase activity of the CobS^m^T complex was the same as that of CobS^m^.

### Data availability

The cryo-EM maps and protein structures generated in this study have been deposited to the Electron Microscopy Data Bank (EMDB) and Protein Data Bank (PDB). The EMDB accession numbers for apo-CobS hexamer are EMD-64243 and EMD-64245. The EMDB accession numbers for the apo-CobS dodecamer are EMD-64244, EMD-64246, and EMD-64247. The EMDB and PDB accession numbers for CobS hexamer without AMPPNP are EMD-64227 and 9UJV. The EMDB and PDB accession numbers for the CobS dodecamer without AMPPNP are EMD-64229 and 9UJX, and EMD-64228 and 9UJW. The EMDB and PDB accession numbers for the CobST with 2 AMPPNPs are EMD-64238 and 9UKG. The EMDB and PDB accession numbers for the apo-CobST are EMD-64242 and 9UKK. The EMDB and PDB accession numbers for the CobST with 3 AMPPNPs are EMD-64239 and 9UKH. The EMDB and PDB accession numbers for the CobS^m^-ATP complex are EMD-64071 and 9UDK. The EMDB and PDB accession numbers for the CobS^m^T-ATP complex are EMD-64235 and 9UKD, and EMD-64237 and 9UKF. The EMDB and PDB accession numbers for the CobS^m^-CobT-shaft-ATP complex are EMD-64236 and 9UKE. The EMDB and PDB accession numbers for the CobST-AMPPNP complex are EMD-64248 and 9UKL, and EMD-64240 and 9UKI. The EMDB and PDB accession numbers for the CobSTN-AMPPNP complex are EMD-64234 and 9UKC, EMD-64233 and 9UKB, and EMD-64232 and 9UKA. The EMDB and PDB accession numbers for the CobN are EMD-64231 and 9UK9. The PDB accession number for CobT_CΔ_ and the CobS_S_-CobT_N_ complex are 21VT and 21VU. This paper does not report any original code.

## Acknowledgements

We thank the staffs from beamlines BL18U1 and BL19U1 of National Facility for Protein Science in Shanghai (NFPSS) at Shanghai Synchrotron Radiation Facility for assistance in data collection and processing. This work was supported by the National Natural Science Foundation of China grants 32471323 (L.L.) and 32371270 (X.C.), and the Anhui Province Outstanding Youth Fund 2308085Y21 (X.C.).

## Author contributions

Experiments were conceived and designed by Y.Z., X.C. and L.L. Protein purifications were done by Y.Z., H.Y., Y.W. and Jian W. Cryo-EM sample preparation and data collection were done by Y.Z., H.C. and L.Y. Crystallization and diffraction data collection were done by Y.Z., M.W., X.W., Jia W. and C.H. High-resolution reconstruction and model building were performed by Y.Z., X.C. and L.L. Figure design, manuscript writing and editing were done by Y.Z., X.C. and L.L. Project supervision was provided by X.C. and L.L. Funding was provided by X.C. and L.L.

## Conflict of interest

The authors declare that they have no conflict of interest.

## Supporting information

**Fig. S1.**
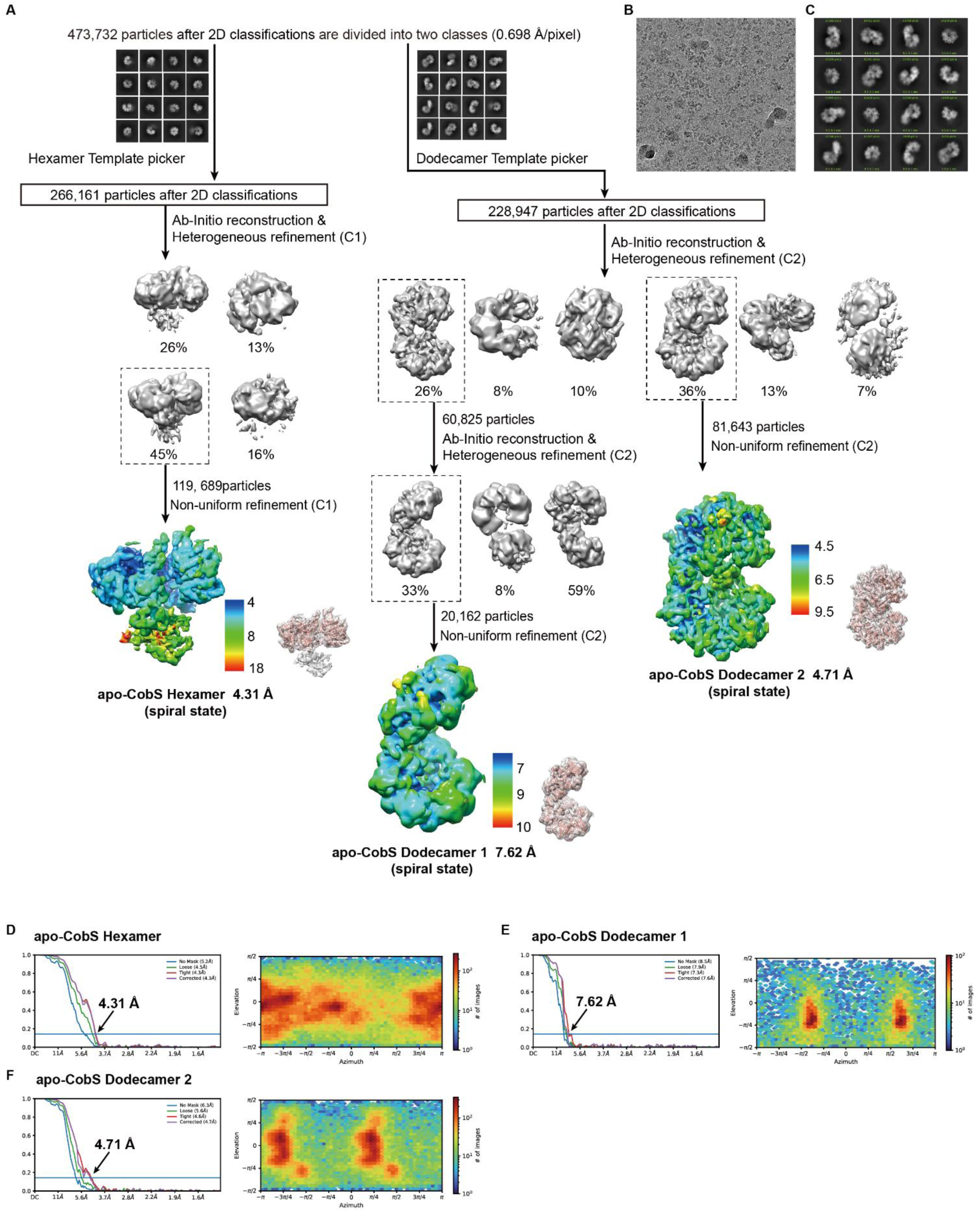
Cryo-EM data processing workflow for apo-CobS. **(A)** Flow chart for the cryo-EM data processing. The maps of local resolution estimations by ResMap with scale bar, and angular distributions of all particles used for the final three-dimensional reconstruction are shown. **(B-C)** Representative cryo-EM micrograph and 2D classes of different views**. (D-F)** FSC curves and angular distributions of apo-CobS maps.

**Fig. S2.**
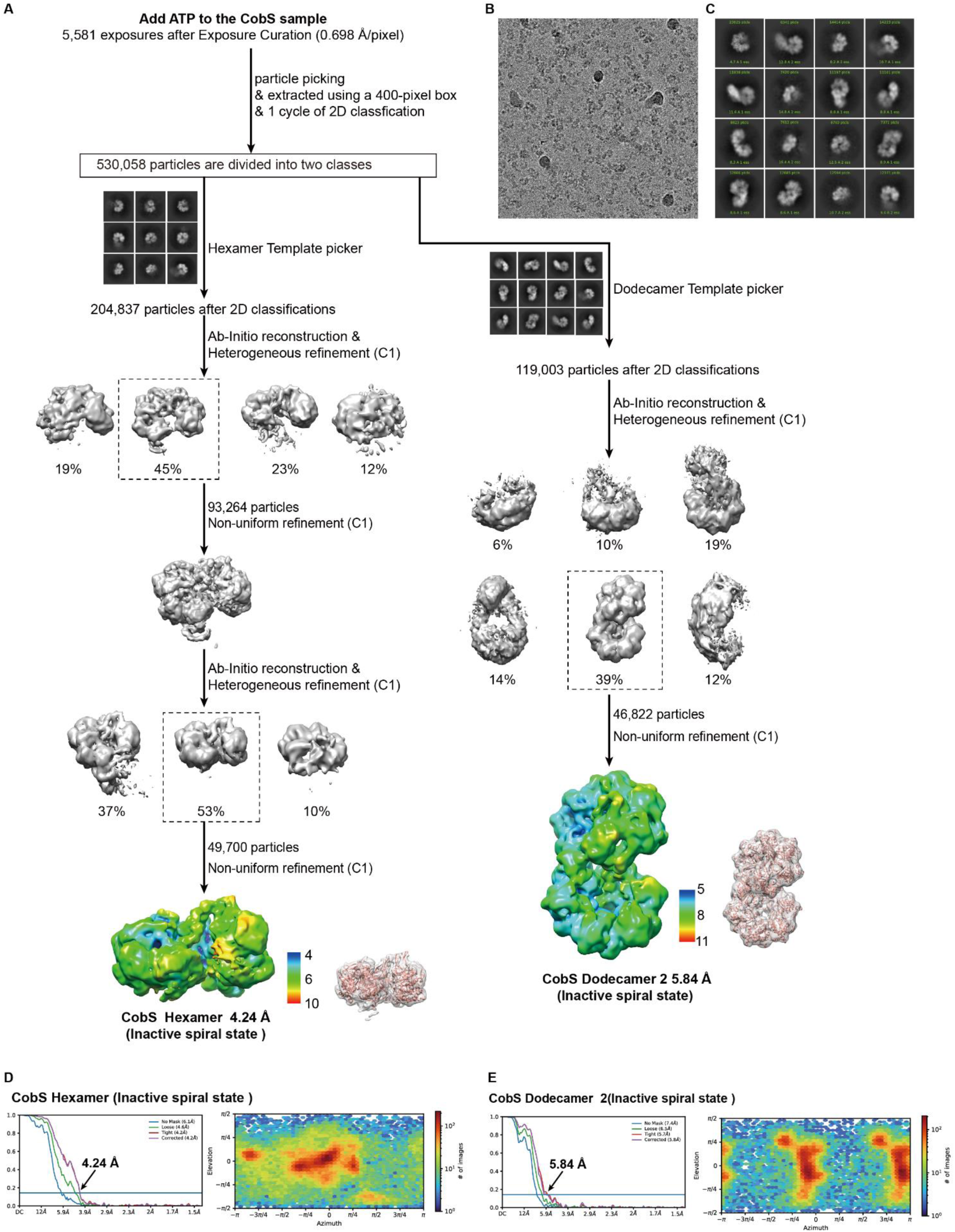
Cryo-EM data processing workflow for CobS with the addition of ATP. **(A)** Flow chart for the cryo-EM data processing. The maps of local resolution estimations by ResMap with scale bar, and angular distributions of all particles used for the final three-dimensional reconstruction are shown. **(B-C)** Representative cryo-EM micrograph and 2D classes of different views. **(D-E)** FSC curves and angular distributions of CobS maps.

**Fig. S3.**
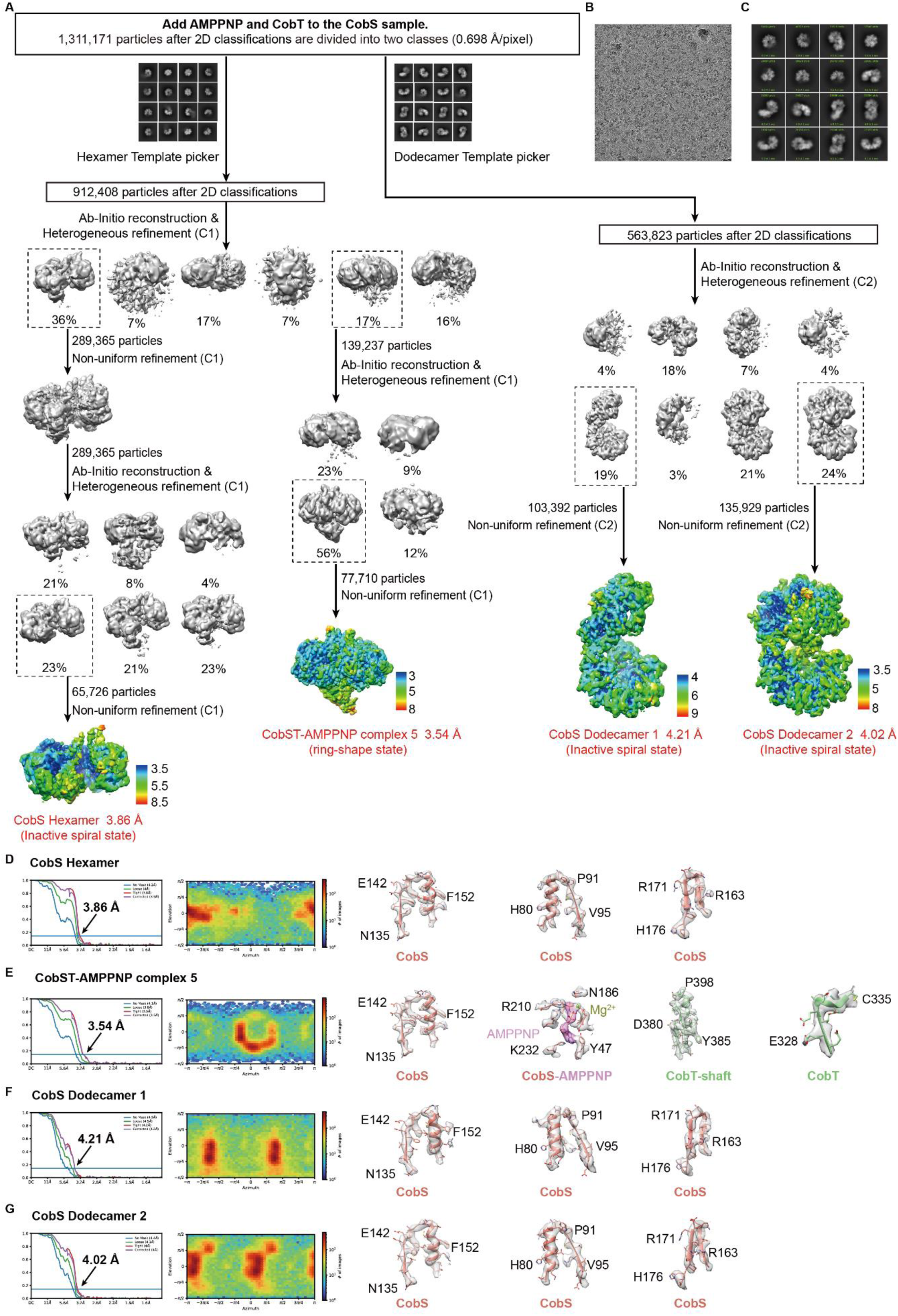
Cryo-EM data processing workflow for CobS with the addition of AMPPNP. **(A)** Flow chart for the cryo-EM data processing. The sample contained trace contaminant CobT. The maps of local resolution estimations by ResMap with scale bar, and angular distributions of all particles used for the final three-dimensional reconstruction are shown. **(B-C)** Representative cryo-EM micrograph and 2D classes of different views. **(D-G)** FSC curves, angular distributions, and representative densities of CobS maps.

**Fig. S4.**
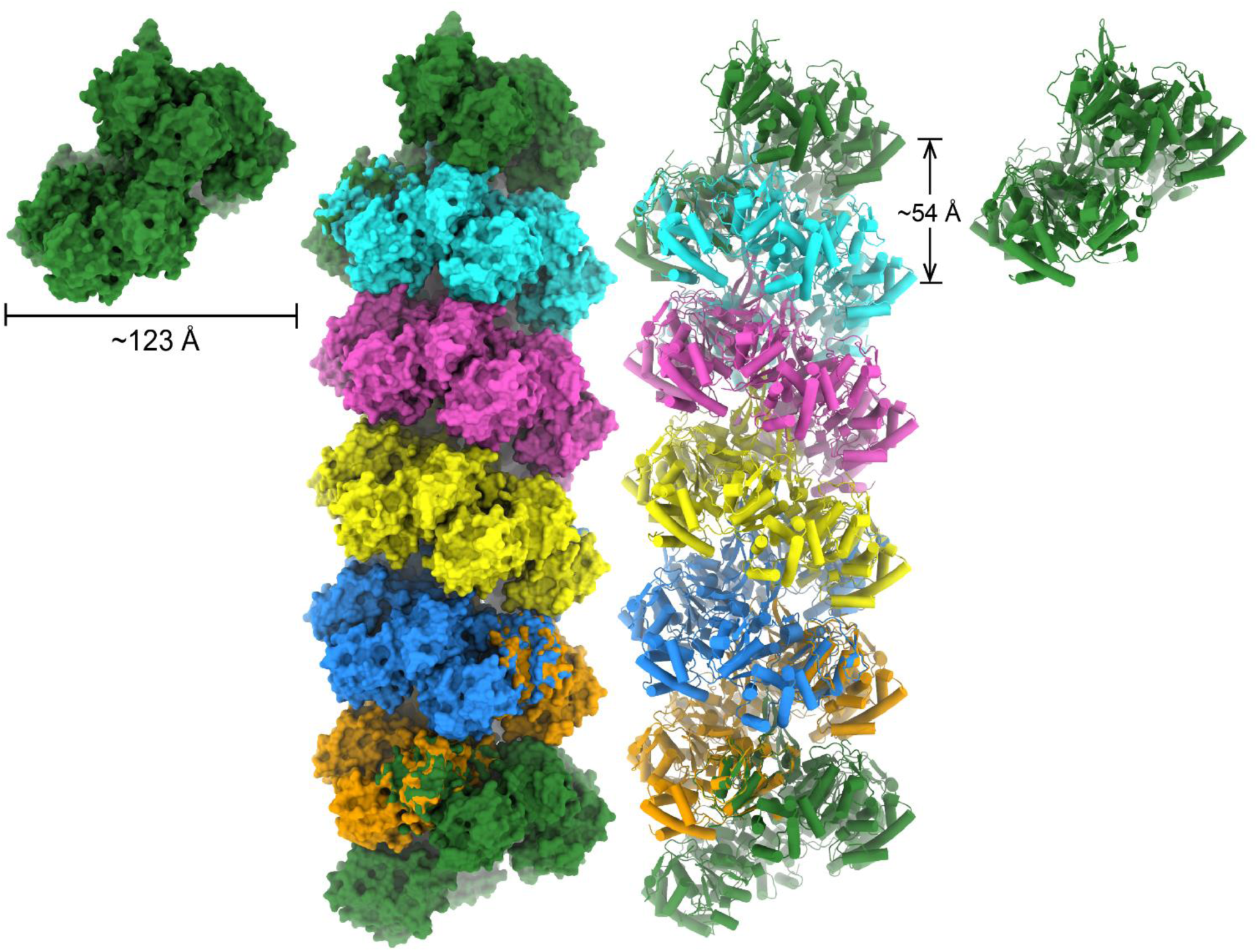
Model of apo-CobS polymer. The first CobSL of one apo-CobS hexamers is superimposed with the last CobSL of its proceeding CobS hexamer. A tubular shape CobS filament is assembled.

**Fig. S5.**
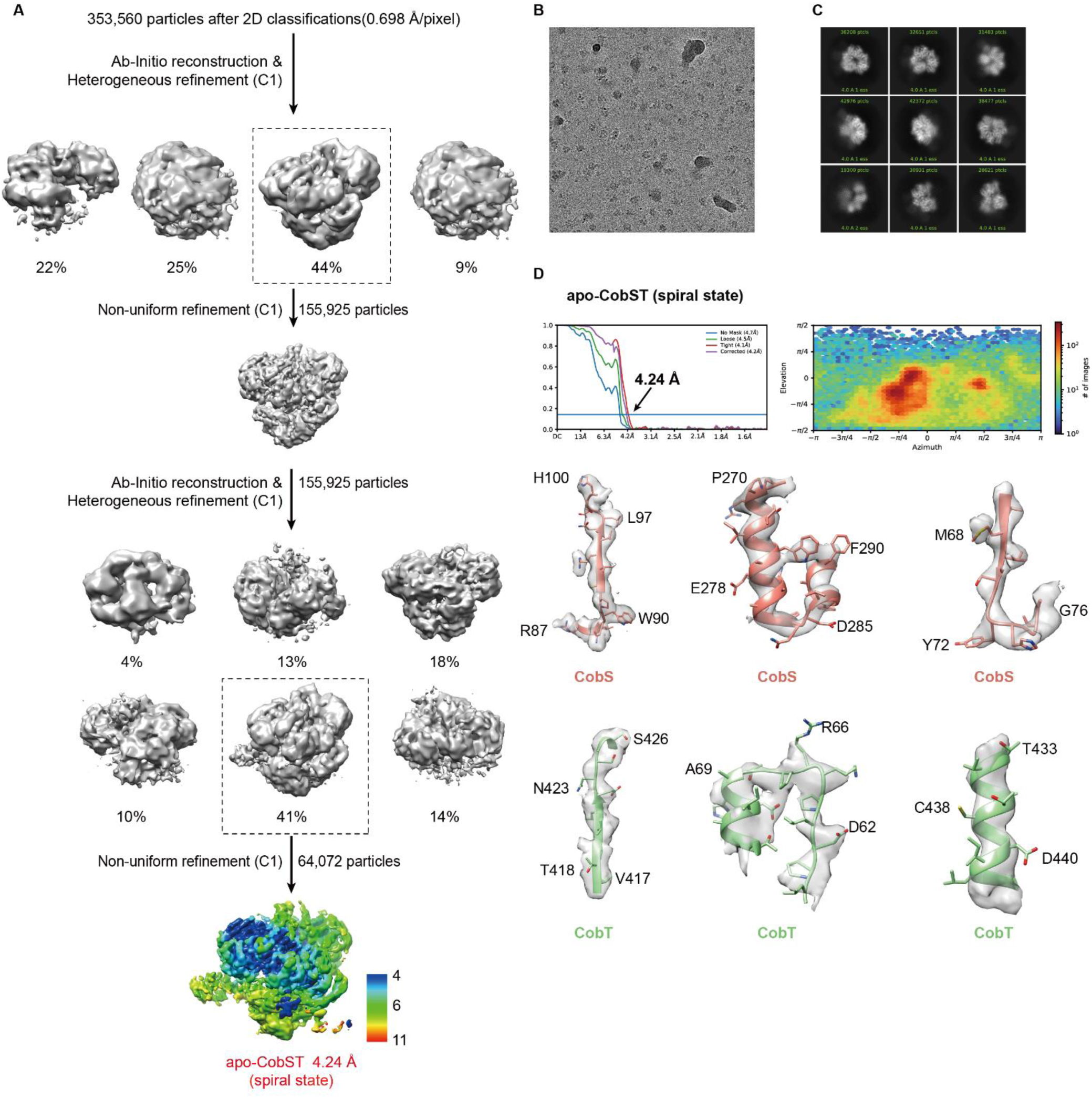
Cryo-EM data processing workflow for apo-CobST. **(A)** Flow chart for the cryo-EM data processing. The maps of local resolution estimations by ResMap with scale bar, and angular distributions of all particles used for the final three-dimensional reconstruction are shown. **(B-C)** Representative cryo-EM micrograph and 2D classes of different views. **(D)** FSC curves, angular distributions, and representative densities of apo-CobST maps.

**Fig. S6.**
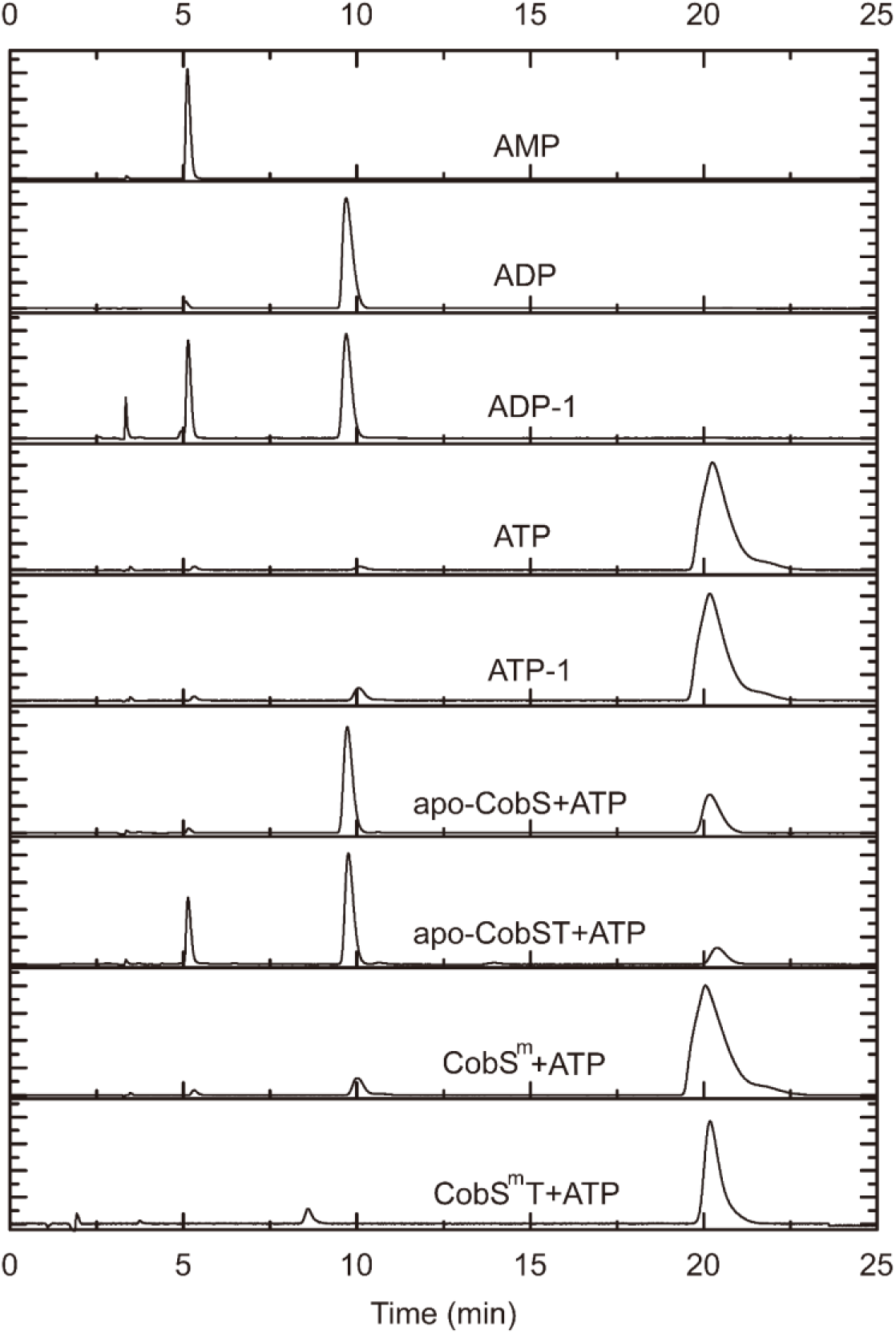
ATP hydrolase assays of CobS, CobST, CobS^m^, and CobS^m^T. The nucleotide AMP, ADP, ADP-1, ATP, ATP-1 were standards.

**Fig. S7.**
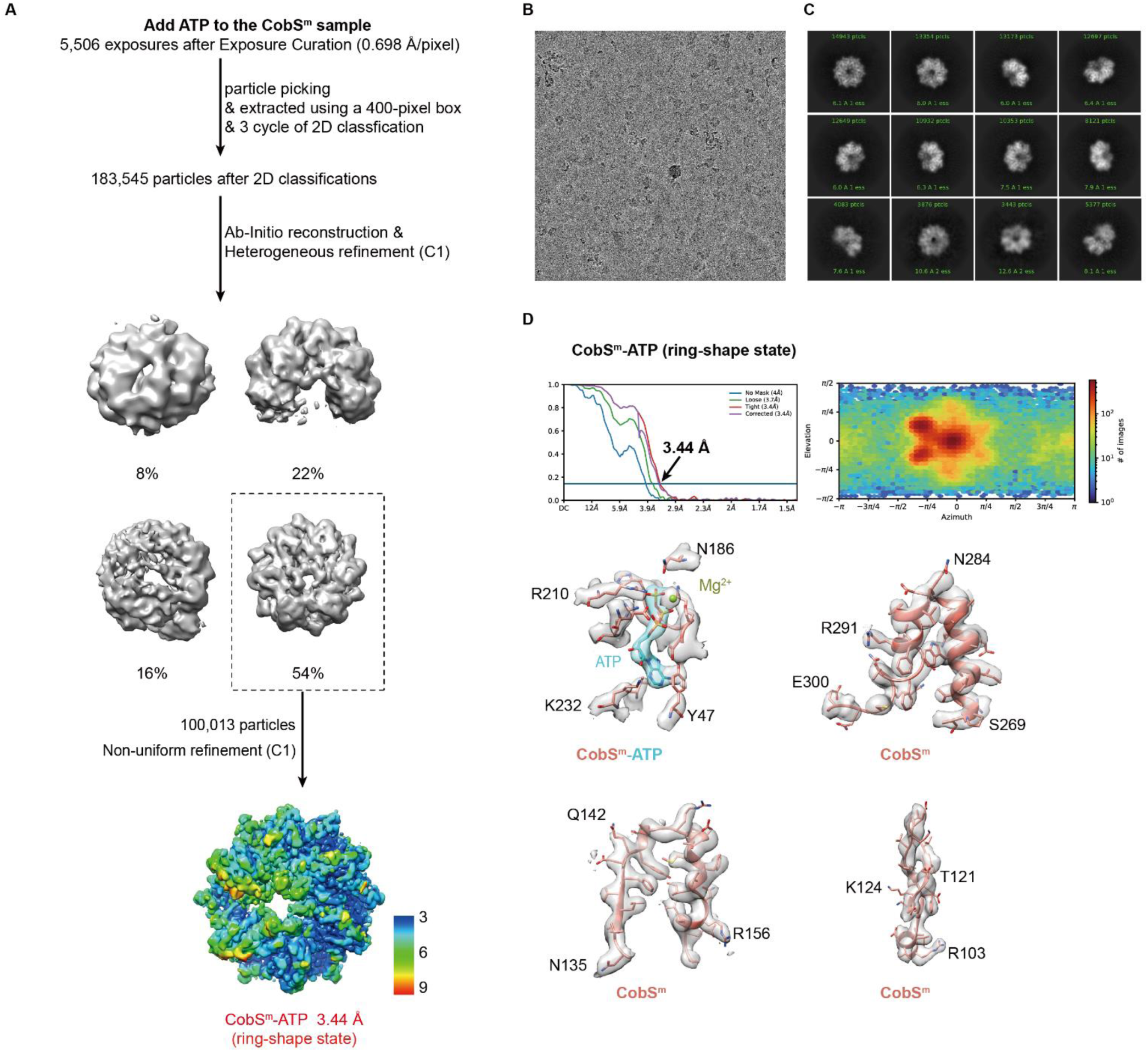
Cryo-EM data processing workflow for CobSm-ATP. **(A)** Flow chart for the cryo-EM data processing. The maps of local resolution estimations by ResMap with scale bar, and angular distributions of all particles used for the final three-dimensional reconstruction are shown. **(B-C)** Representative cryo-EM micrograph and 2D classes of different views. **(D)** FSC curves, angular distributions, and representative densities of CobSm-ATP maps.

**Fig. S8.**
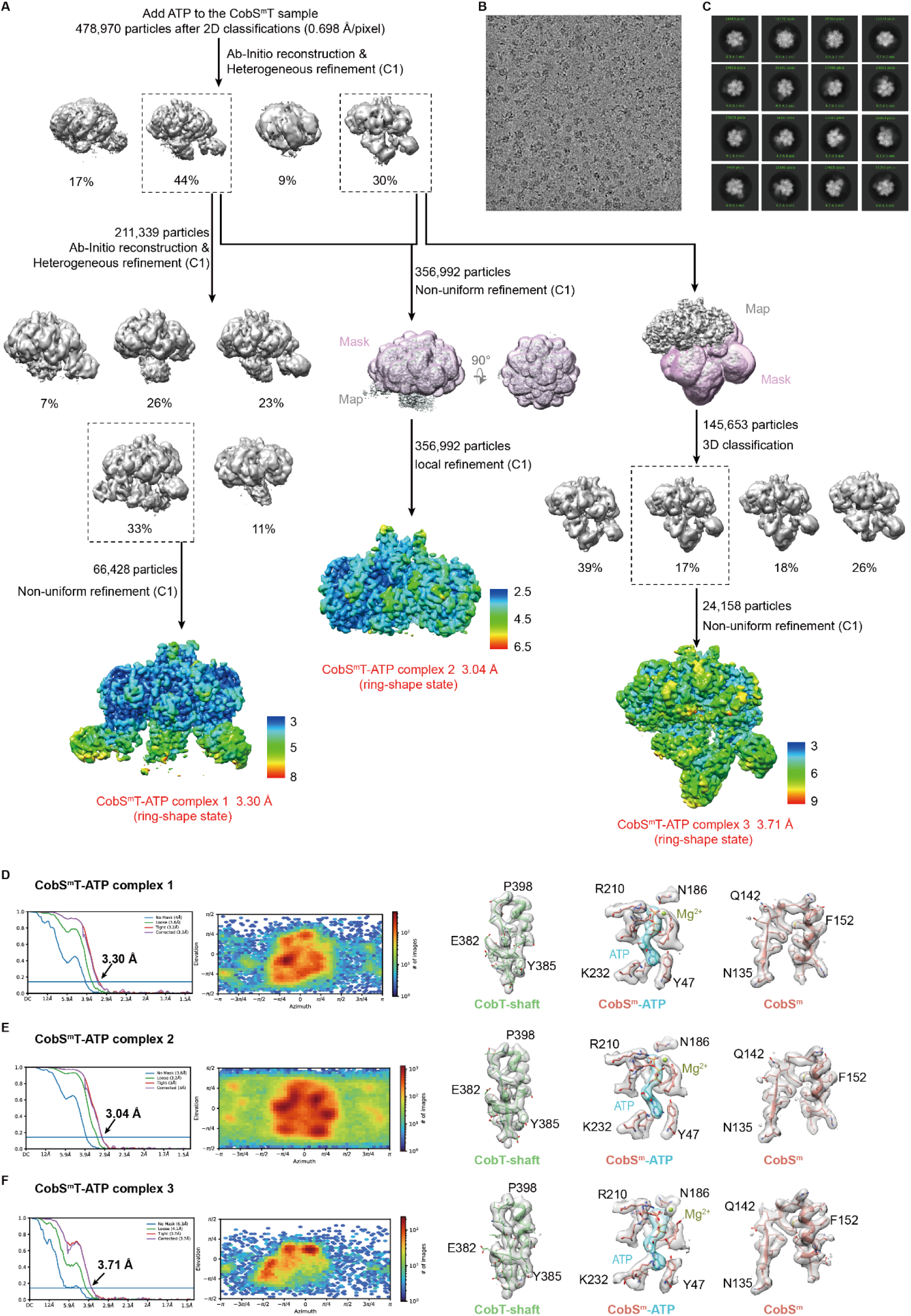
Cryo-EM data processing workflow for CobSmT-ATP. **(A)** Flow chart for the cryo-EM data processing. The maps of local resolution estimations by ResMap with scale bar, and angular distributions of all particles used for the final three-dimensional reconstruction are shown. **(B-C)** Representative cryo-EM micrograph and 2D classes of different views. **(D-F)** FSC curves, angular distributions, and representative densities of CobSmT-ATP maps.

**Fig. S9.**
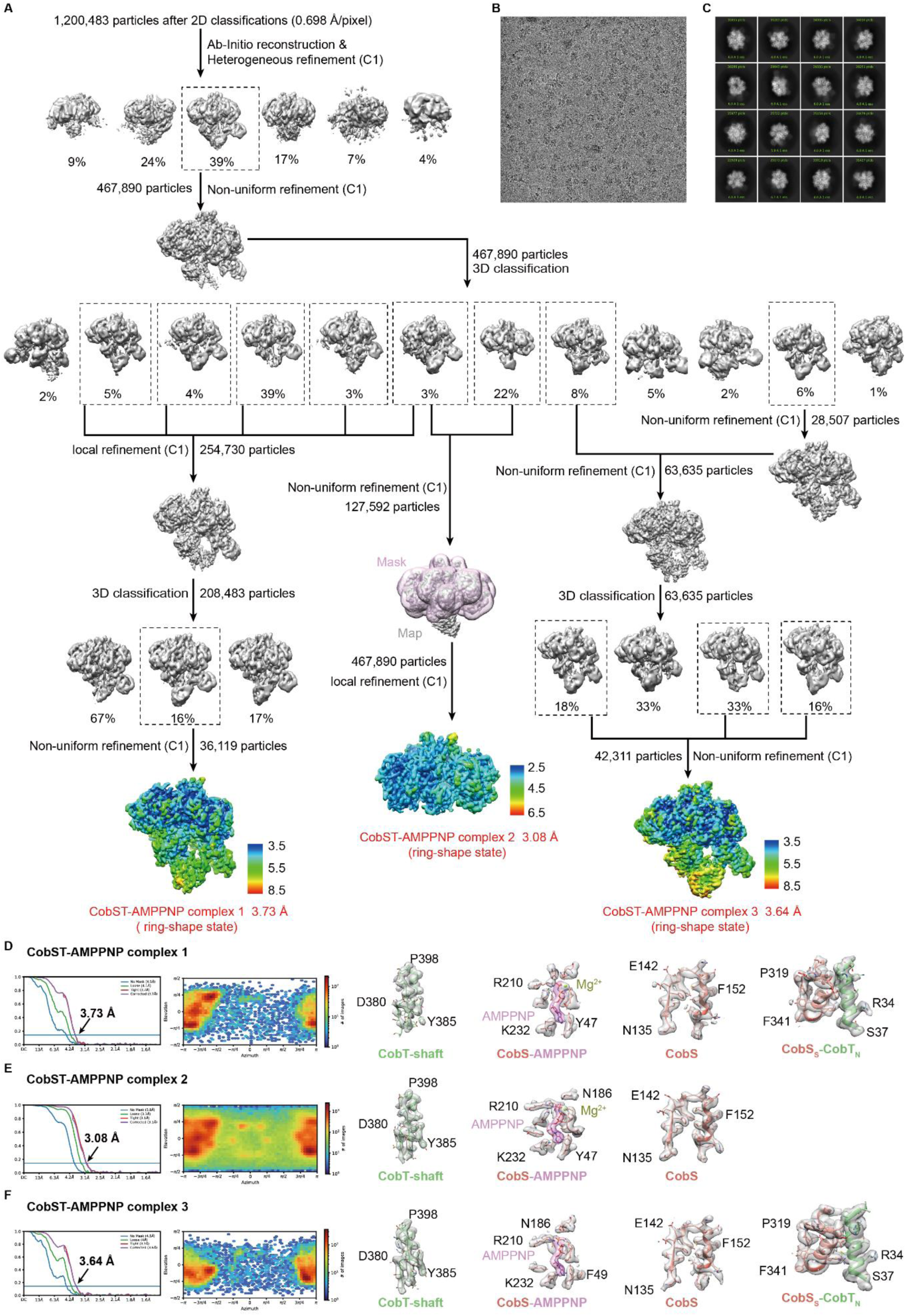
Cryo-EM data processing workflow for CobST-AMPPNP. **(A)** Flow chart for the cryo-EM data processing. The maps of local resolution estimations by ResMap with scale bar, and angular distributions of all particles used for the final three-dimensional reconstruction are shown. **(B-C)** Representative cryo-EM micrograph and 2D classes of different views. **(D-F)** FSC curves, angular distributions, and representative densities of CobST-AMPPNP maps.

**Fig. S10.**
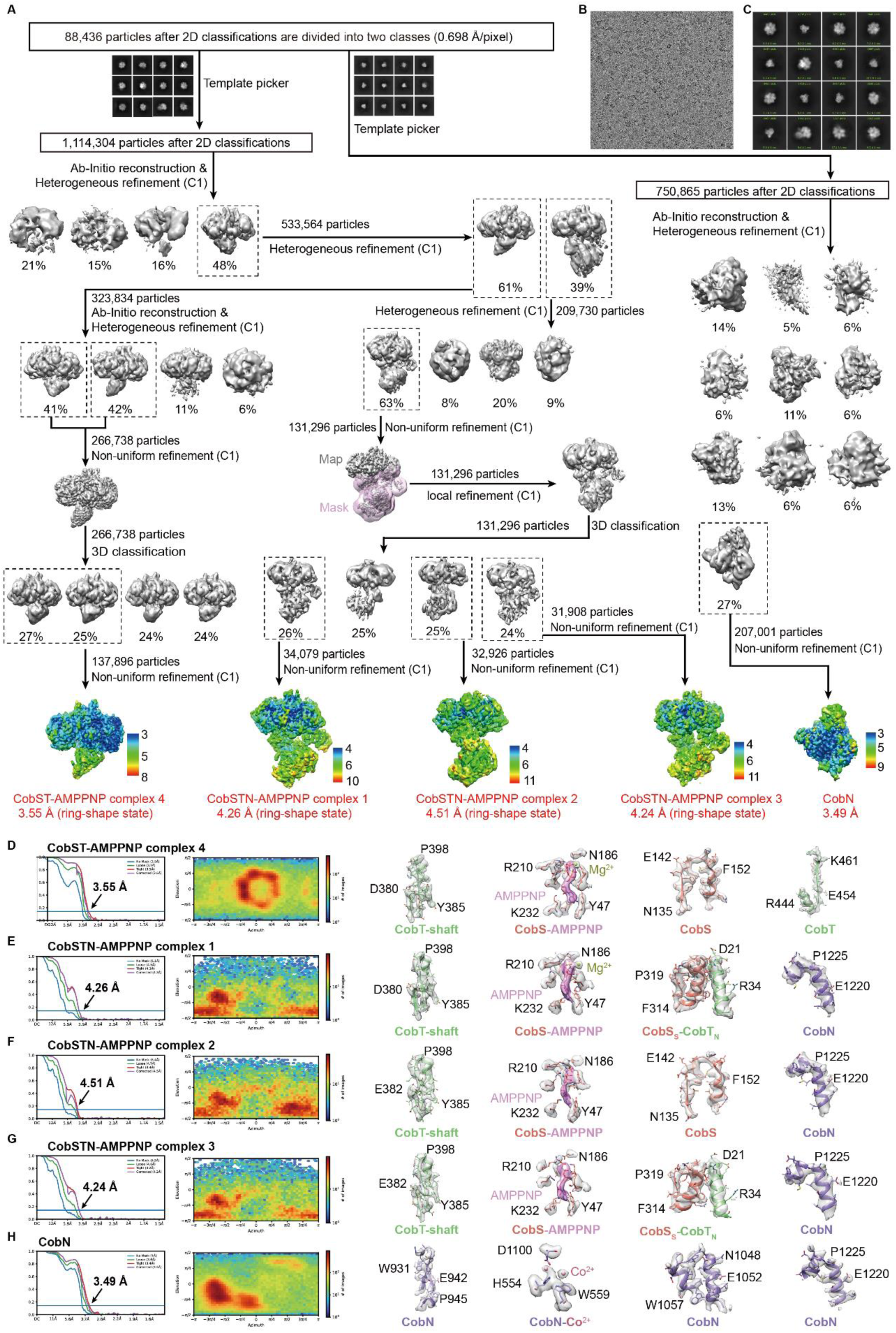
Cryo-EM data processing workflow for CobSTN-AMPPNP. **(A)** Flow chart for the cryo-EM data processing. The maps of local resolution estimations by ResMap with scale bar, and angular distributions of all particles used for the final three-dimensional reconstruction are shown. **(B-C)** Representative cryo-EM micrograph and 2D classes of different views. **(D-H)** FSC curves, angular distributions, and representative densities of CobSTN-AMPPNP map

**Fig. S11.**
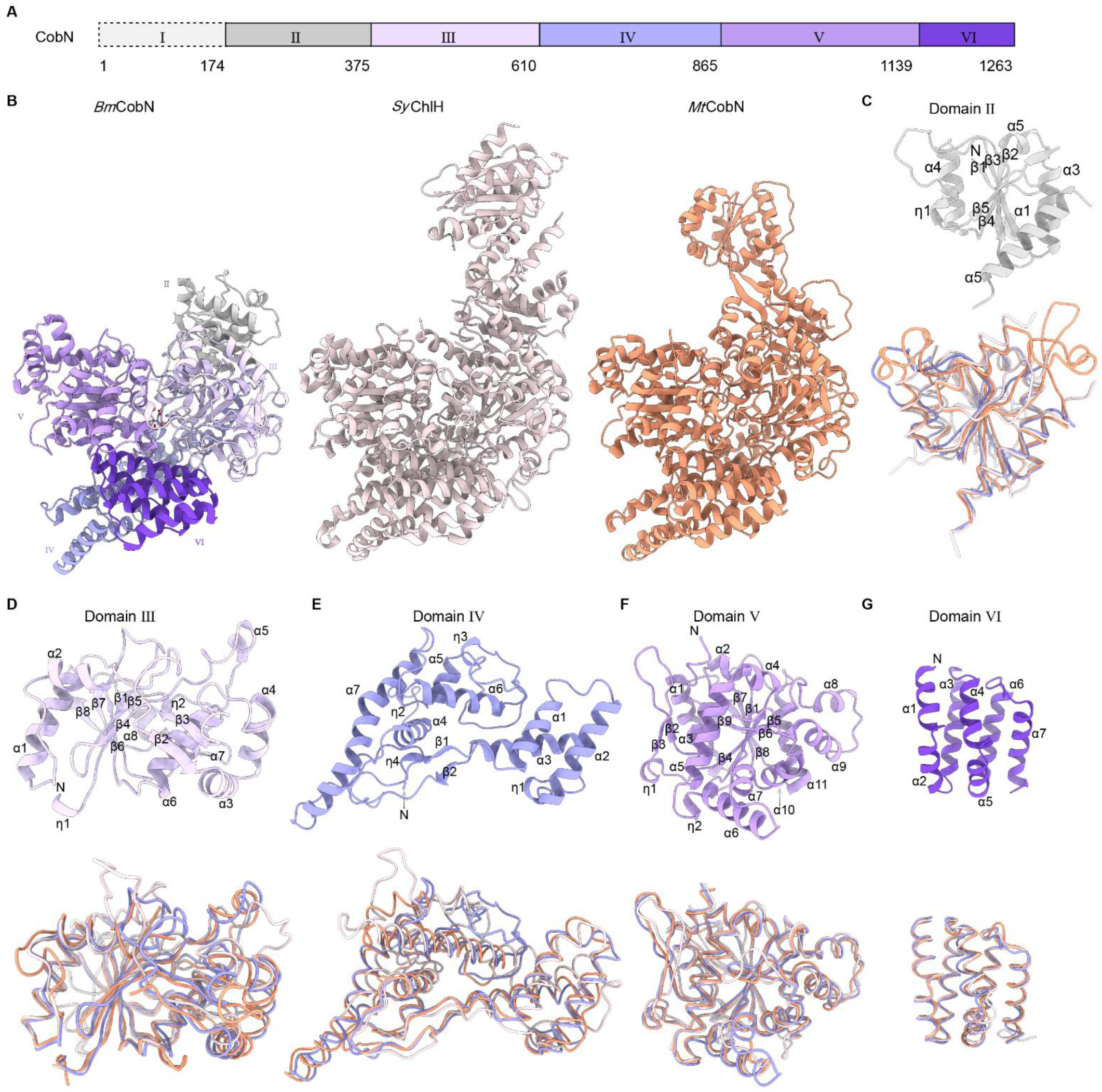
Comparison of BmCobN, SyChlH, and MtCobN structures. **(A)** Overall structures of BmCobN (colored same as in Fig. 5B), SyChlH (light pink) and MtCobN (dark salmon). **(B-G)** Domains II-VI of BmCobN, shown in cartoon with the secondary structures numbered (up); and comparison with its corresponding domains of MtCobN and SyChlH, shown in ribbon (down).

**Table S1.** Cryo-EM data collection of apo-CobS and the CobS with the additional of ATP.

|  | apo-CobS<br>Hexamer<br>(EMDB-64245) | apo-CobS<br>Dodecamer 1<br>(EMDB-64246) | apo-CobS<br>Dodecamer 2<br>(EMDB-64247) | CobS<br>Hexamer<br>(EMDB-64243) | CobS<br>Dodecamer 2<br>(EMDB-64244) |
| --- | --- | --- | --- | --- | --- |
| Magnification | 165,000 | 165,000 | 165,000 | 165,000 | 165,000 |
| Voltage (kV) | 200 | 200 | 200 | 200 | 200 |
| Electron dose (e<sup>-</sup>/Å<sup>2</sup>) | 50 | 50 | 50 | 50 | 50 |
| Defocus range (μm) | -0.4 to -2 | -0.4 to -2 | -0.4 to -2 | -0.4 to -2 | -0.4 to -2 |
| Pixel size (Å/pixel) | 0.698 | 0.698 | 0.698 | 0.698 | 0.698 |
| Symmetry imposed | C1 | C2 | C2 | C1 | C1 |
| Initial particle image (no.) | 266,161 | 228,947 | 228,947 | 204,837 | 119,003 |
| Final particle image (no.) | 65,726 | 20,162 | 81,643 | 49,700 | 46,822 |
| Map resolution (Å) |  |  |  |  |  |
| 0.143 FSC threshold | 4.31 | 7.62 | 4.71 | 4.24 | 5.84 |

**Table S2.**
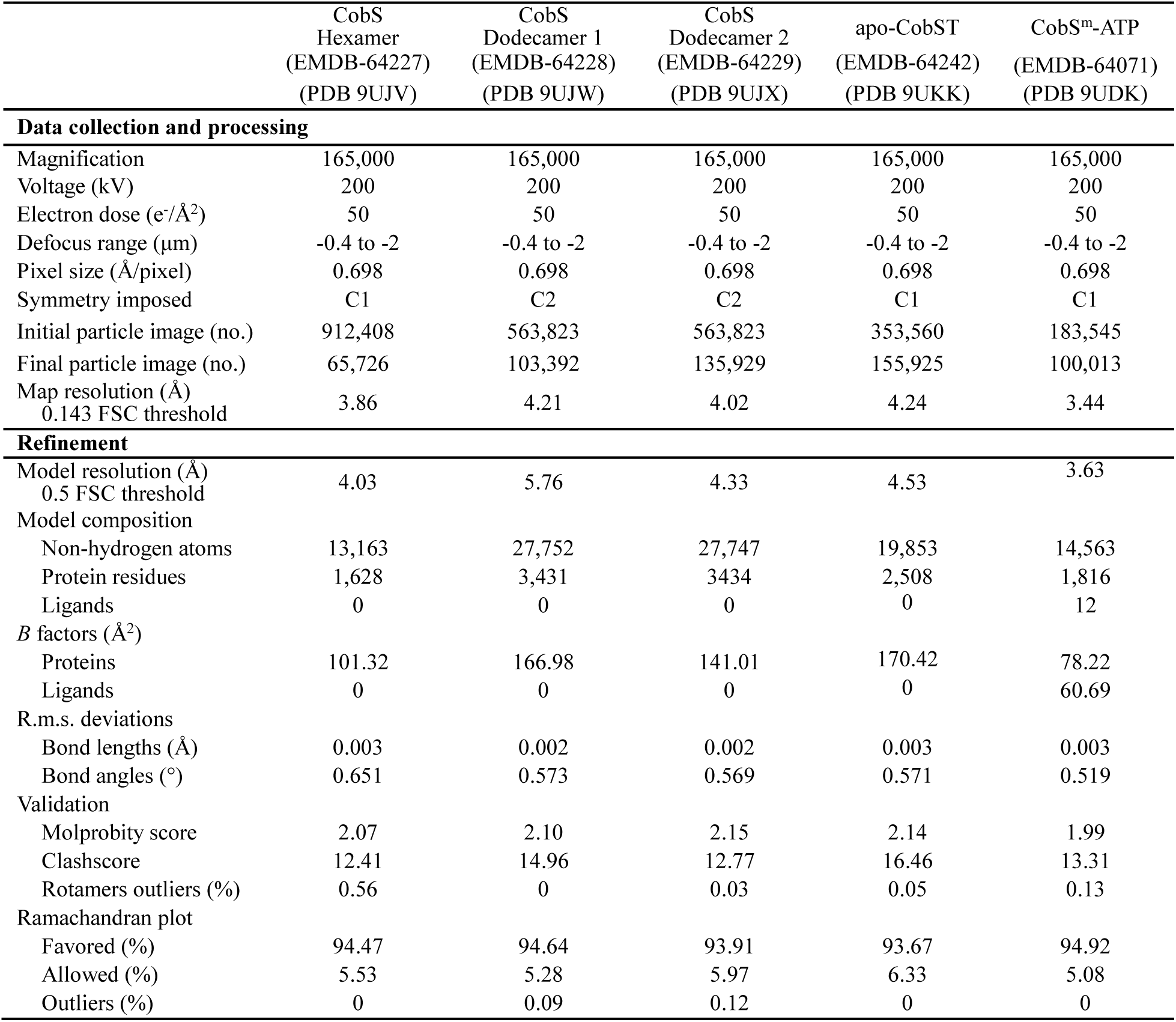
Cryo-EM data collection, refinement and validation statistics of the CobS with the additional of AMPPNP sample, apo-CobST and the CobS^m^-ATP complex.

**Table S3.**
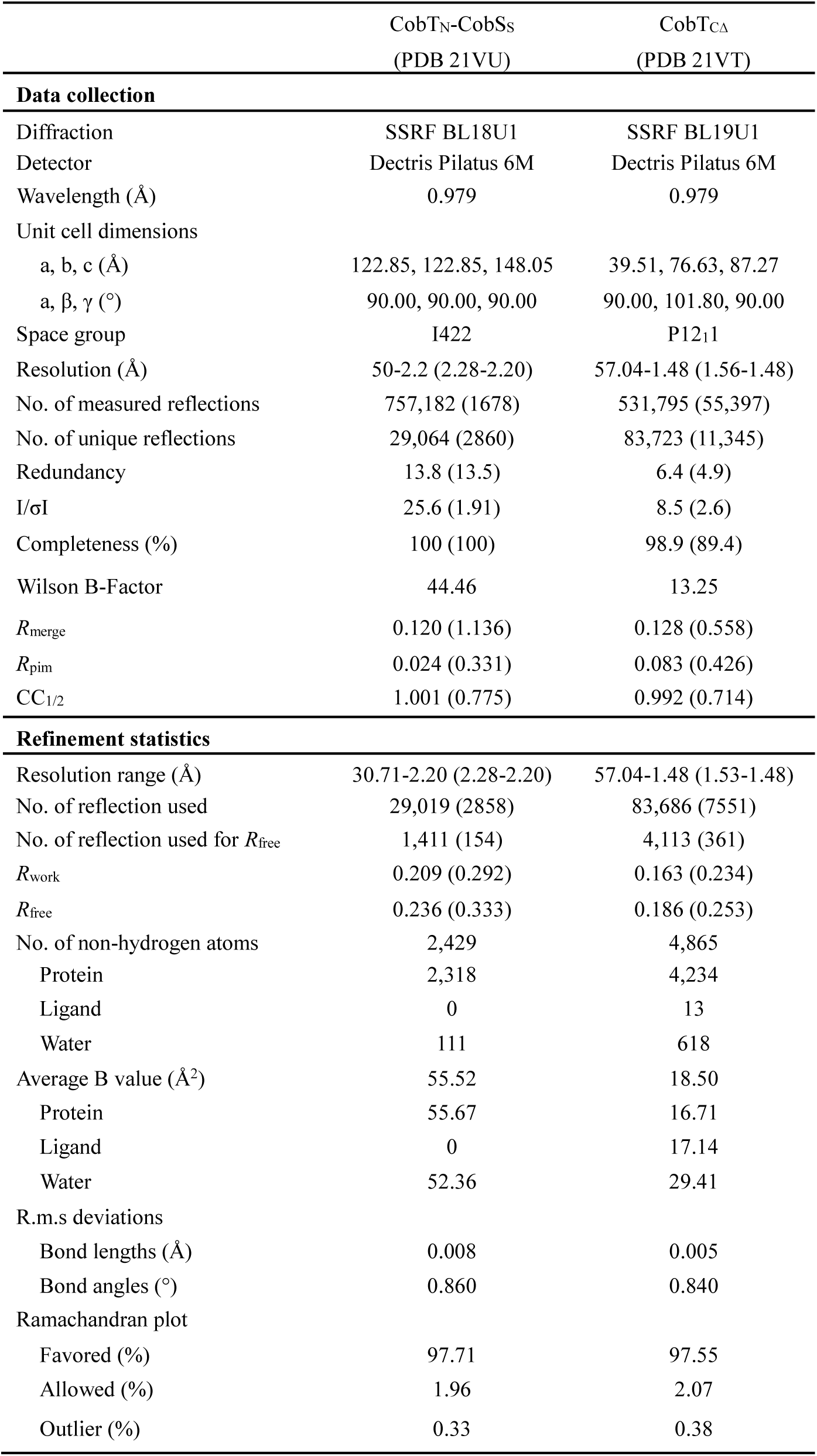
Crystal structure data collection and refinement statistics of CobT_CΔ_ and CobT_N_-CobS_S_ complex.

**Table S4.** Cryo-EM data collection, refinement and validation statistics of CobS^m^T-ATP complex.

|  | CobS<sup>m</sup>T-ATP<br>complex 1<br>(EMDB-64237)<br>(PDB 9UKF) | CobS<sup>m</sup>T-ATP<br>complex 2<br>(EMDB-64236)<br>(PDB 9UKE) | CobS<sup>m</sup>T-ATP<br>complex 3<br>(EMDB-64235)<br>(PDB 9UKD) |
| --- | --- | --- | --- |
| **Data collection and processing** |  |  |  |
| Magnification | 165,000 | 165,000 | 165,000 |
| Voltage (kV) | 200 | 200 | 200 |
| Electron dose (e<sup>-</sup>/Å<sup>2</sup>) | 50 | 50 | 50 |
| Defocus range (μm) | -0.4 to -2 | -0.4 to -2 | -0.4 to -2 |
| Pixel size (Å/pixel) | 0.698 | 0.698 | 0.698 |
| Symmetry imposed | C1 | C1 | C1 |
| Initial particle image (no.) | 478,970 | 478,970 | 478,970 |
| Final particle image (no.) | 66,428 | 356,997 | 24,158 |
| Map resolution (Å) | 3.30 | 3.04 | 3.71 |
| 0.143 FSC threshold |  |  |  |
| **Refinement** |  |  |  |
| Model resolution | 3.38 | 3.23 | 3.94 |
| 0.5 FSC threshold |  |  |  |
| Model composition |  |  |  |
| Non-hydrogen atoms | 23,302 | 15,199 | 21,255 |
| Protein residues | 2,577 | 1,898 | 2,673 |
| Ligands | 11 | 11 | 10 |
| *B* factors (Å<sup>2</sup>) |  |  |  |
| Protein | 87.11 | 71.38 | 89.91 |
| Ligand | 66.17 | 67.50 | 74.62 |
| R.m.s. deviationa |  |  |  |
| Bonds length (Å) | 0.004 | 0.004 | 0.004 |
| Bonds angles (°) | 0.578 | 0.484 | 0.640 |
| Validation |  |  |  |
| Molprobity score | 1.86 | 1.86 | 2.28 |
| Clashscore | 11.89 | 9.12 | 17.69 |
| Rotamers outliers (%) | 0.57 | 0.19 | 0.45 |
| Ramachandran plot |  |  |  |
| Favored (%) | 94.68 | 94.52 | 90.42 |
| Allowed (%) | 5.32 | 5.48 | 9.58 |
| Outliers (%) | 0 | 0 | 0 |

**Table S5.** Cryo-EM data collection, refinement and validation statistics of CobST-AMPPNP complex.

|  | CobST-AMPPNP<br>complex 1<br>(EMDB-64248)<br>(PDB 9UKL) | CobST-AMPPNP<br>complex 2<br>(EMDB-64241)<br>(PDB 9UKJ) | CobST-AMPPNP<br>complex 3<br>(EMDB-64240)<br>(PDB 9UKI) |
| --- | --- | --- | --- |
| **Data collection and processing** |  |  |  |
| Magnification | 165,000 | 165,000 | 165,000 |
| Voltage (kV) | 200 | 200 | 200 |
| Electron dose (e<sup>-</sup>/Å<sup>2</sup>) | 50 | 50 | 50 |
| Defocus range (μm) | 0.4 to 2 | 0.4 to 2 | 0.4 to 2 |
| Pixel size (Å/pixel) | 0.698 | 0.698 | 0.698 |
| Symmetry imposed | C1 | C1 | C1 |
| Initial particle image (no.) | 1,200,483 | 1,200,483 | 1,200,483 |
| Final particle image (no.) | 32,948 | 254,730 | 42,311 |
| Map resolution (Å) | 3.73 | 3.08 | 3.64 |
| 0.143 FSC threshold |  |  |  |
| **Refinement** |  |  |  |
| Model resolution (Å) | 3.94 | 3.27 | 3.96 |
| 0.5 FSC threshold |  |  |  |
| Model composition |  |  |  |
| Non-hydrogen atoms | 20,995 | 15,003 | 20,759 |
| Protein residues | 2,649 | 1,883 | 2,618 |
| Ligands | 6 | 6 | 6 |
| *B* factors (Å<sup>2</sup>) |  |  |  |
| Protein | 102.33 | 76.92 | 104.61 |
| Ligand | 82.72 | 71.41 | 75.78 |
| R.m.s.deviation |  |  |  |
| Bonds lengths (Å) | 0.003 | 0.004 | 0.003 |
| Bonds angles (°) | 0.640 | 0.534 | 0.584 |
| Validation |  |  |  |
| Molprobtity score | 2.19 | 1.87 | 2.06 |
| Clashscore | 17.41 | 8.75 | 14.06 |
| Rotamers Outliers (%) | 0.45 | 0.37 | 0.36 |
| Ramachandran plot |  |  |  |
| Favored (%) | 92.87 | 94.10 | 93.98 |
| Allowed (%) | 7.13 | 5.90 | 6.02 |
| Outliers (%) | 0 | 0 | 0 |

**Table S6.** Cryo-EM data collection, refinement and validation statistics of CobSTN-AMPPNP complex and CobN.

|  | CobSTN-AMPPNP complex 1<br>(EMDB-64234)<br>(PDB 9UKC) | CobSTN-AMPPNP complex 2<br>(EMDB-64233)<br>(PDB 9UKB) | CobSTN-AMPPNP complex 3<br>(EMDB-64232)<br>(PDB 9UKA) | CobN<br>(EMDB-64231)<br>(PDB 9UK9) |
| --- | --- | --- | --- | --- |
| **Data collection and processing** |  |  |  |  |
| Magnification | 165,000 | 165,000 | 165,000 | 165,000 |
| Voltage (kV) | 200 | 200 | 200 | 200 |
| Electron dose (e<sup>-</sup>/Å<sup>2</sup>) | 50 | 50 | 50 | 50 |
| Defocus range (μm) | -0.4 to -2 | -0.4 to -2 | -0.4 to -2 | -0.4 to -2 |
| Pixel size (Å/pixel) | 0.698 | 0.698 | 0.698 | 0.698 |
| Symmetry imposed | C1 | C1 | C1 | C1 |
| Initial particle image (no.) | 1,114,304 | 1,114,304 | 1,114,304 | 750,865 |
| Final particle image (no.) | 34,079 | 32,926 | 31,908 | 207,001 |
| Map resolution (Å)<br>0.143 FSC threshold | 4.26 | 4.51 | 4.24 | 3.49 |
| **Refinement** |  |  |  |  |
| Model resolution (Å)<br>0.5 FSC threshold | 4.54 | 6.24 | 4.52 | 4.05 |
| Model composition |  |  |  |  |
| Non-hydrogen atoms | 25,970 | 23,905 | 25,824 | 7,837 |
| Protein residues | 3,299 | 3,032 | 3,282 | 1,023 |
| Ligands | 6 | 3 | 3 | 2 |
| *B* factors (Å<sup>2</sup>) |  |  |  |  |
| Protein | 141.19 | 164.29 | 159.00 | 83.32 |
| Ligand | 109.84 | 124.51 | 115.63 | 90.08 |
| R.m.s. deviation |  |  |  |  |
| Bonds length (Å) | 0.003 | 0.003 | 0.003 | 0.004 |
| Bonds angles (°) | 0.627 | 0.608 | 0.599 | 0.707 |
| Validation |  |  |  |  |
| Molprobity score | 2.18 | 2.20 | 2.17 | 2.08 |
| Clashscore | 16.92 | 19.16 | 16.89 | 16.06 |
| Rotamers outliers (%) | 0 | 0.04 | 0 | 0.13 |
| Ramachandran plot |  |  |  |  |
| Favored (%) | 92.82 | 93.55 | 93.25 | 94.60 |
| Allowed (%) | 7.18 | 6.45 | 6.75 | 5.4 |
| Outliers (%) | 0 | 0 | 0 | 0 |

## Notes

### Competing Interest Statement

The authors have declared no competing interest.

